# Dynamic Supercoiling Sponsors Transcription Amplification by MYC

**DOI:** 10.64898/2026.01.19.700398

**Authors:** Rajiv Kumar Jha, Fedor Kouzine, Bo Wang, James D. Phelan, Subhendu K Das, Brian A Lewis, David Levens

## Abstract

MYC dysregulation occurs in most cancers provoking, profound changes in gene expression. Although sometimes considered to be an E-box-dependent transcription factor, an alternate model posits MYC to be a universal amplifier of active genes. Although MYC is associated with accelerated pause release, the full extent of its participation in the transcription cycle remains poorly illuminated. MYC also stimulates topoisomerases to resolve topological issues that complicate DNA transactions; whether and how this stimulation is coordinated with or independent of its transcriptional role has not been examined. We have developed a genetic tool that discriminates between MYC’s ability to activate versus amplify reporter gene expression in any cell. This system enables the interrogation of genes, cofactors, and compounds that execute or modulate MYC activity. Using a combination of biochemical and cellular assays, we reveal the dynamic interplay between MYC-driven transcription amplification and DNA supercoiling. The early stages of transcription are highly sensitive to the level of DNA supercoiling. Reducing the activity of TOP1 either through genetic knockdown or low-dose inhibition potentiates transcription amplification by MYC. This enhancement is associated with pre-initiation complex (PIC) stabilization by DNA supercoiling. MYC helps to mobilize these stabilized PICs.

**Significance Statement:** MYC is dysregulated in most cancers, yet its precise functions in transcription are poorly understood. Using a genetic system that distinguishes transcriptional activation from amplification, we reveal a functional interplay between MYC-driven amplification and DNA supercoiling. Reduced topoisomerase I activity increases DNA-supercoiling that in turn stabilizes PICs to provide more substrate for MYC-dependent transcription amplification. Thus, DNA topology as a critical regulator of MYC-dependent transcription.

## Introduction

The MYC protooncogene has been generally considered to be a transcription factor acting at physiological levels to activate a broad panel of target genes upon binding to their promoter E-boxes, a DNA element with the sequence CACGTG (1, 2). At pathologically elevated oncogenic levels, MYC has been shown to bind promoters and enhancers promiscuously to exaggerate target gene expression (3, 4). An alternative interpretation is that MYC is incapable of initiating target gene expression but amplifies the expression of already active genes according to their existing transcriptional activity (1, 3, 5–7). In this scenario, E-boxes influence, but do not determine promoter output. Constitutive or induced pathways that activate the appropriate transcription factors provide the input signal(s) that determine amplifier output.

The mechanisms of MYC action, whether as an activator or as an amplifier, have been incompletely elucidated. Engaging a large and complex set of partner proteins, MYC may help to deliver transcription regulatory molecules to promoters and enhancers via binding to E-boxes or interacting with the transcription machinery at different stages of the transcription cycle (1, 8–10). Whether the delivery of these molecules is part of a coordinated reaction scheme or MYC simply deposits cargo into a transcription condensate thereby increasing the local concentration of necessary components is unknown. That MYC can activate or inhibit the enzymatic activities of some partners may argue for some degree of coordination. For example, MYC forms a discrete complex that includes both topoisomerases 1 and 2 in which their catalytic activities are dramatically enhanced and in the case of TOP1 reprogrammed—in the absence of MYC, TOP1 fails to engage DNA until after pause release whereas in its presence TOP1 forms covalent complexes at transcription start sites (TSS) (11–13). Topoisomerases help to tune the topology and conformation of template DNA to favor initiation, pausing or elongation. During the earliest stages of transcription, preservation of negative supercoiling at start sites would assist in pre-initiation complex assembly (14–17). But this supercoiling must then be overcome to allow the template-engaged RNA polymerase II (RNAPII) to advance beyond the start site, from initiation through early elongation, pausing, and thereafter pause release and elongation (18, 19). The firing of promoters in both prokaryotes and eukaryotes occurs in bursts of activity directing the synthesis of several transcripts, punctuated by more prolonged periods of inactivity. To amplify transcription, MYC extends the “on-times” but not the frequency of transcriptional bursts. The build-up and release of torsional stress within the DNA template have been shown to be a major determinate of dynamics of transcriptional bursting (20–22). Empirically, both MYC and topoisomerases control transcription bursting, and MYC directly regulates topoisomerase activity. So, the regulation of topoisomerase activity may help to adjust the gain of MYC amplified transcription but may also alter the frequency of promoter firing (23, 24). Here we explore the coordination, integration, and the synergistic or antagonistic actions of topoisomerases with MYC on a model gene and across the genome.

## Results

### Establishing a cell-based tool to study MYC-mediated transcriptional regulation

Whether MYC is a conventional transcription factor that switches genes on, or an amplifier operating in the service of other transcription factors, is still a matter of some controversy. Because of the complex arrangements of binding sites, redundant regulatory features, and chromatin architecture that might modify MYC’s action at natural gene promoters, we sought to develop a system to illuminate the essential action(s) of MYC in gene expression (Fig. 1A). This system included reading frames encoding: (1) a MYC-Estrogen receptor fusion expressed from the strong EF1-α promoter (25–27); this constitutively expressed protein almost instantly translocates to nuclei upon the addition of tamoxifen. (2) The Tet-On 3G protein comprised of Tet-on DNA binding domain with three minimal VP16 activation domains expressed from the strong constitutive hPGK promoter (25); upon treatment of cells with doxycycline (Dox) this synthetic transcription factor is immediately activated to express mCherry fluorescent protein from the TRE3GS promoter. (3) This promoter includes multiple Tet operators that dock Tet-VP16 and a TATA-box to direct initiation; binding sites for all other transcription factors have been eliminated. Another variant of this reporter includes an E-box inserted between the TATA-box and the Tet operators (Fig. 1A). The utilization of mCherry provides a marker for flow cytometry and cell sorting as well as enabling direct visualization of expression in individual cells in tissue culture. (4) Lyt-2/CD8a driven by the SV40 early promoter provides a constitutive cell surface reporter to mark cells carrying this system in the absence of mCherry expression. Because these components were embedded in a lentivirus backbone, they were insulated by the viral LTRs from the influence of nearby genomic regulatory elements that might otherwise confound analysis. This lentivirus (with or without the E-box at TRE3GS promoter) was stably transduced into immortalized, non-transformed MCF10A breast epithelial cells.

**Fig. 1.**
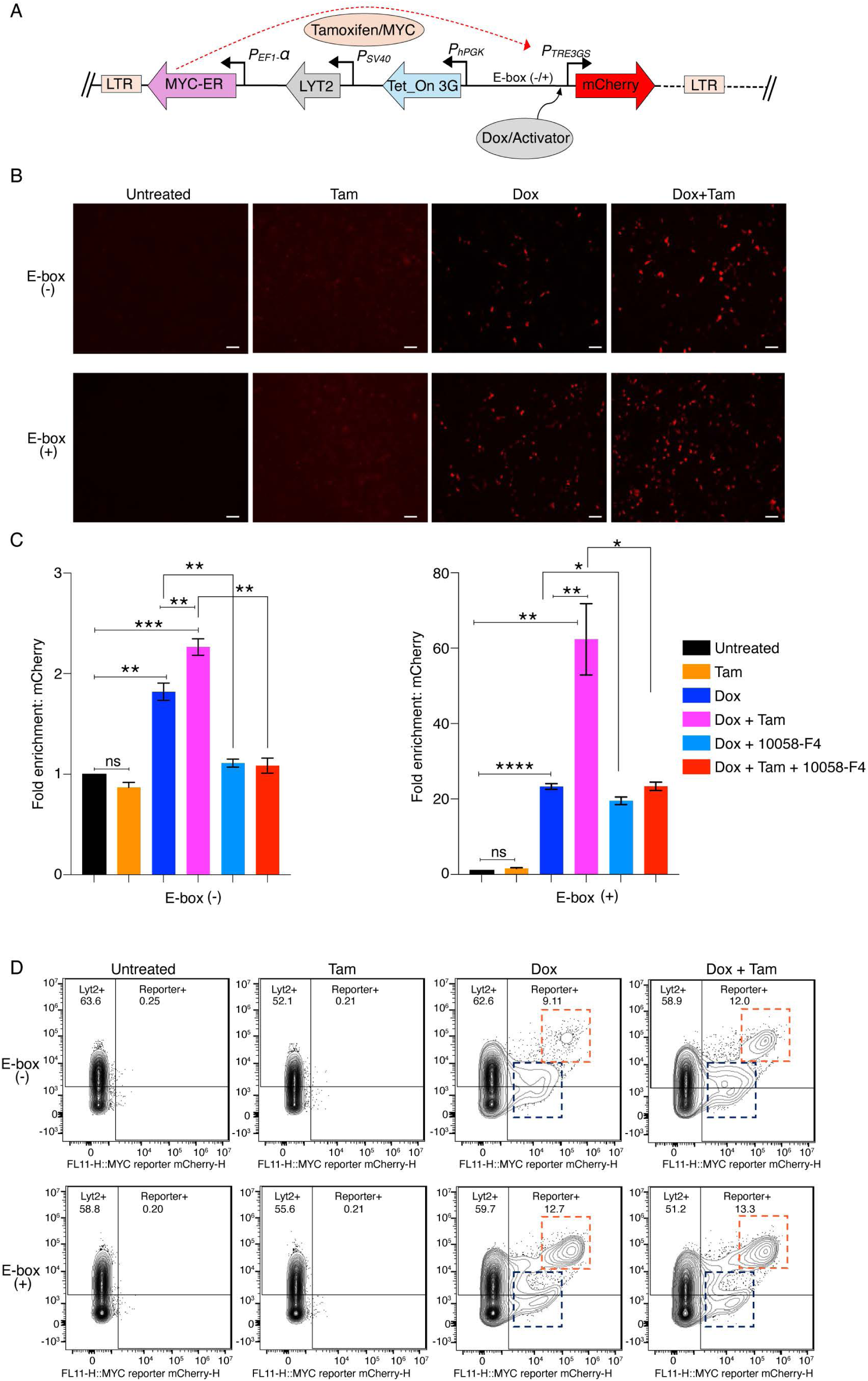
A cell-based assay establishes MYC as a transcription amplifier. *(A)* Schematic of the lentiviral constructs used to generate the inducible MYC-ER system to assess role of MYC in transcription. Transduced cells harbor a tamoxifen (Tam)-responsive MYC-ER fusion, a Tet-On 3G transactivator, and a doxycycline (Dox)-inducible mCherry reporter. Upon tamoxifen treatment, MYC-ER translocates to the nucleus, and its effect on transcription monitored by doxycycline-driven mCherry expression. *(B)* MCF10A cells expressing lentiviral constructs (with or without E-box) in the mCherry promoter were treated with or without Dox (150 ng/ml) and Tam (200 µM). Representative microscopic images from at least three independent experiments are shown (scale bars, 200 µm). *(C)* Quantification of fold change in mCherry mRNA expression in MCF10A cells expressing lentiviral construct without (left) and with (right) E-box under the indicated Dox, Tam conditions and ± MYC inhibitor 10058-F4 (calculated value ± SEM). For all statistical analyses (two-way ANOVA): *, **, and ***, indicate P < 0.05, 0.01,and 0.001, respectively; ns denotes nonsignificant. *(D)* MCF10A cells expressing lentiviral mCherry constructs with or without an E-box in the promoter were cultured under indicated Dox (150 ng/ml) and Tam (200 µM) conditions and analyzed by flow cytometry. Cells were analyzed by flow cytometry and mCherry fluorescence is shown on the X-axis, and LYT2 was detected using an anti-CD8α antibody on the Y-axis. Data were gated on live singlet cells gated for mCherry and LYT2 (Alexa-647) channels; transcriptionally activated and amplified populations are highlighted with blue and red dashed boxes, respectively. Data represents from at least three independent experiments.

To test MYC’s action as a conventional transcription factor operating through a promoter proximal E-box, E-box (+) or E-box (-), MCF10A cells were treated with tamoxifen and examined microscopically for mCherry fluorescence. The complete absence of fluorescent cells without doxycycline and tamoxifen treatments confirmed the very low output of the unactivated TRE3GS promoter (Fig. 1B). Tamoxifen treatment alone failed to elicit any increase in mCherry signal (Fig. 1B). In contrast, a large fraction of cells fluoresced following doxycycline treatment, confirming the functionality of TRE3GS-activated promoter and the mCherry reporter. Notably, mCherry output plateaued with saturating levels of doxycycline.

Changes in mCherry fluorescence were faithfully paralleled at the RNA level as assessed using RTqPCR, confirming that system controls mCherry mRNA expression as intended (Fig. 1C).

### MYC amplifies the transcriptional output of promoters activated by transcription factors

Cells were either treated with or without the activator (doxycycline) to induce mCherry expression and compared in the absence and presence of tamoxifen to activate MYC-ER. Cells treated with tamoxifen in the absence of the activator did not express mCherry regardless of the presence or absence of a promoter E-box (Fig. 1B); thus, MYC alone could not activate reporter transcription. But when cells were treated with the activator to induce mCherry expression, expression was further elevated by switching MYC-ER on with tamoxifen (Fig. 1B). To further validate and better quantify these results, cells were analyzed by flow-cytometry under these same experimental conditions. Cells were gated on the basis of mCherry and LYT2 expression. Lacking Tet-On 3G activation, mCherry-positive cells were absent irrespective of tamoxifen treatment or the presence of a promoter E-box (Fig. 1D). When cells were treated with doxycycline, mCherry-positive populations were detected, and their fluorescence was intensified with tamoxifen treatment (Fig. 1D). The mCherry-positive population observed with the doxycycline alone represents transcription activation by Tet-On 3G (dashed blue box), while the additional increase in fluorescence intensity (dashed red box) induced by MYC-ER reflects transcription amplification (Fig. 1D). Because MYC-ER also augments LYT2 expression, the activated and amplified mCherry-positive populations were cleanly resolved. The high mCherry/high-LYT2 zone defines the population of cells hosting MYC-amplified transcription. That the E-box (+) promoter yielded both activated and amplified populations even in the absence of tamoxifen (Fig. 1D) likely reflected E-box assisted recruitment of the endogenous MYC to sponsor a low amount of amplification (Fig 1B). Notably, though the E-box increased the number of cells in the amplified zone, it did not intensify the degree of mCherry fluorescence relative to the non-E-box reporter. Turning on MYC-ER with tamoxifen further increased the amplified population (Fig. 1D). To test if transcription amplification required MYC to dimerize with MAX, MCF10A cells were treated with 10058-F4, a widely used inhibitor of MYC-MAX heterodimerization (28) and monitored by flow cytometry. Indeed, 10058-F4 depressed the amplified mCherry population (SI Appendix, Fig. S1A). To check the robustness of the system, E-box (+) and E-box (-) lentiviruses were transduced into OCI-LY1 lymphoma cells that express endogenous MYC at much higher levels than MCF10A cells (SI Appendix, Fig. S1B). Treatment of OCI-LY1 cells with increasing concentrations of 10058-F4 progressively reduced transcription amplification (SI Appendix, Fig. S1C). However, approximately five-fold higher concentrations of drug were required to achieve the same level of inhibition as in MCF10A cells, indicating that effective inhibitor concentration scaled up with the elevated level of MYC (SI Appendix, Fig. S1C). The improved and more potent MYC-MAX heterodimerization inhibitors MYCi971 and MYCi361(29) each reduced the population of amplified mCherry cells (SI Appendix, Fig. S1D) at lower doses than 10058-F4 in accord with their higher affinities for MYC-MAX. The negative impact of MYC dimerization inhibitor 10058-F4 on reporter mCherry mRNA levels was confirmed using RTqPCR (Fig. 1C). These data demonstrate that MYC-MAX functions as a transcription amplifier. Thus, this robust cell-based system is a good platform to investigate the molecular mechanisms and factors governing MYC-mediated transcriptional amplification.

### Inhibition of TOP1 enhances transcription amplification by MYC, independent of DNA damage

Transcription progresses through multiple stages during which the molecular machinery and the chromatin-embedded DNA template are progressively remodeled and modified. It is still not known whether the transcription-cycle proceeds through a largely invariant, stereotypical set of reactions, or whether different combinations of complexes and factors alter the reaction pathway at different genes according to the state of the cell or stochastic fluctuations. It has been argued that to amplify transcription globally, MYC must interact with multiple components at different stages of the transcription cycle (1). Regardless of the precise mechanism, increased transcription will pump more topological stress into active DNA templates and their embracing chromatin fibers (16, 30–35). Unless mitigated, this stress would impede further MYC-driven transcription. MYC-topoisomes (MYC-TOP1-TOP2A) have been proposed to resolve the topological issues provoked by high levels of transcription (13); however, the functional interplay between MYC’s action at promoters during transcription amplification with topoisomes has not been explored. To interrogate the relationship of DNA supercoiling and transcription amplification, we examined the influence of DNA topoisomerase inhibitors on doxycycline-activated mCherry expression in the presence or absence of MYC-ER. The topoisomerase inhibitors included camptothecin (CPT), a selective inhibitor of Topoisomerase I (TOP1), as well as etoposide and ICRF-187, that each target Topoisomerase II (TOP2), but with different modes of action (36–38).

Inclusion of a low level of CPT (0.5 μM) with doxycycline treatment dramatically increased the number of cells in the amplified zone (Fig. 2A). In the absence of doxycycline, CPT neither activated nor boosted mCherry (SI Appendix, Fig. S2A). Thus, either CPT amplified transcription in parallel with and independent of MYC, or else it boosted the population of MYC-responsive cells. Inhibiting TOP2 using 20 μM etoposide only slightly increased amplification (∼2-fold increase) whereas 5 μM ICRF-187 had no significant impact on reporter expression (Fig. 2A). To better characterize the influence of CPT on mCherry reporter activation and on transcription amplification, cells were treated across a range of drug concentrations.

**Fig. 2.**
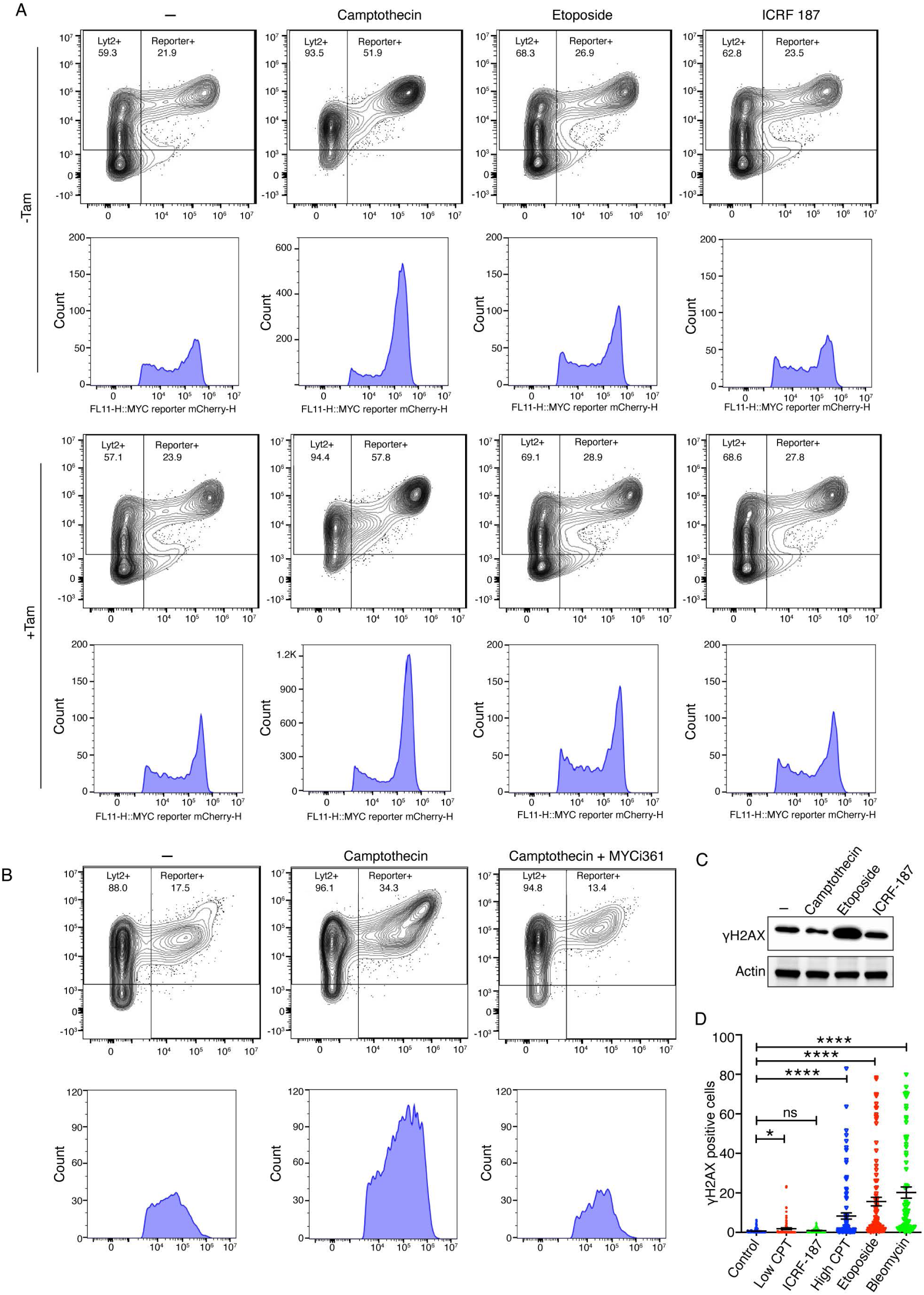
Inhibition of TOP1 enhances transcription amplification by MYC, independent of DNA damage *(A)* MCF10A cells expressing lentiviral E-box (+) mCherry constructs were cultured with Dox (150 ng/mL) ± Tam (200 µM) and treated with the indicated topoisomerase inhibitors—camptothecin (0.5 µM), etoposide (20 µM), or ICRF-187 (5 µM). Cells were analyzed by flow cytometry and mCherry fluorescence is shown on the X-axis, and LYT2 was detected using an anti-CD8α antibody on the Y-axis. Data were gated on live singlet cells gated for mCherry and LYT2 (Alexa-647) channels. Histograms below show the fluorescence intensity distributions of mCherry-positive populations under each condition. *(B)* MCF10A cells expressing lentiviral E-box (-) mCherry constructs were cultured with Dox (150 ng/mL) and treated with indicated combination of CPT (100 nM) and MYCi361 (2.5 µM). Cells were analyzed by flow cytometry and mCherry fluorescence is shown on the X-axis, and LYT2 was detected using an anti-CD8α antibody on the Y-axis. Data were gated on live singlet cells gated for mCherry and LYT2 (Alexa-647) channels. Histograms below show the fluorescence intensity distributions of mCherry-positive populations under each condition. *(C)* Immunoblot showing 𝞬H2AX levels in cells treated with camptothecin (0.5 µM), etoposide (20 µM), or ICRF-187 (5 µM). Actin served as a loading control. *(D)* Immunofluorescence microscopy of MCF10A cells (shown in SI Appendix, Fig. S2E) expressing lentiviral E-box (+) mCherry constructs under treatment with Dox (150 ng/mL) ± Tam (200 µM) and treated with the indicated topoisomerase inhibitors- low (100 nM) versus high dose CPT (2.5 µM), etoposide (20 μM), ICRF-187 (5 μM), and bleomycin (5 μM). Nuclei were stained with DAPI and Nuclear γH2AX was quantified for 100 cells for each treatment (calculated values ± SEM from three biological replicates). Each dot corresponds to γH2AX intensity per nucleus. For all statistical analyses (t-test): *, and ****, indicate P < 0.05, and 0.0001, respectively; ns denotes nonsignificant.

Titration of CPT from 4 nM to 2.5 μM revealed dramatic stimulation of transcription amplification at much lower concentrations than typically used to inhibit topoisomerase activity (SI Appendix, Fig. S2B) (39–44). The increase in transcription amplification plateaued at 100-500 nM CPT and notably declined thereafter (SI Appendix, Fig. S2B). At low stimulatory doses CPT markedly increased the fraction of cells populating the amplified zone while also supporting a small but consistent increase in the fluorescence intensity of the mCherry reporter. The increase in the amplified population following treatment with 100 nM CPT was further confirmed by fluorescence microscopy (SI Appendix, Fig. S2C).

Addition of tamoxifen only modestly augmented the transcription amplification enforced by low-dose CPT (Fig. 2A). The limited stimulation by MYC over CPT suggested that amplification had already plateaued. Perhaps low-dose CPT had primed amplification by the endogenous MYC. To test whether CPT-enforced amplification was dependent upon or independent of MYC action, cells were treated with both CPT and MYC-inhibitors. Notably, in the presence of CPT, amplification remained fully sensitive to MYC inhibition by MYCi361 (Fig. 2B) and by 10058-F4 inhibitor (SI Appendix, Fig. S2D), even without tamoxifen, indicating that CPT acted through endogenous MYC to amplify mCherry expression. As expected, MYC inhibition compromised amplification by MYC-ER. These results suggest that low-level CPT enabled more efficient utilization of the endogenous MYC to augment reporter expression. In fact, when the mCherry promoter harbored an E-box, the endogenous MYC was able to saturate reporter output even prior to the addition of tamoxifen. That TOP1 inhibition by CPT increased reporter expression, suggested that TOP1 might tonically repress transcription *in vivo,* just as it does in transcription reactions *in vitro* (45, 46).

Topoisomerase inhibitors might alter reporter expression by disturbing the generation and transmission of torsional stress within the DNA fiber, or more indirectly by creating DNA damage that interferes with gene expression. CPT and etoposide damage DNA by stabilizing covalent cleavage complexes between DNA and TOP1 or TOP2, respectively (38, 47, 48). The TOP1- and TOP2- covalent cleavage complexes in turn provoke a series of events that alter global gene expression, potentially complicating the separation of the transcriptional changes that result directly from the inhibition of topoisomerase activity from those reflecting the DNA damage. Therefore, γ-H2AX was monitored as a surrogate for DNA damage (49) following treatment with low CPT, etoposide, or ICRF-187. Global changes in γ-H2AX levels were assessed by immunoblot. Low concentrations of CPT and ICRF-187 did not appreciably elevate global γ-H2AX levels compared to control cells, whereas an increase was clearly noted following etoposide treatment (Fig. 2C). Apparently, most TOP1 cleavage complexes trapped at low levels of CPT are transient and reversible, and so do not provoke a DNA-damage response. To directly visualize intracellular foci of DNA damage, cells were examined by fluorescence microscopy after staining for γ-H2AX. In contrast to high dose CPT, etoposide and bleomycin [DNA-damaging control (50)] which stained almost all cells, most cells with low dose CPT or ICRF187 remained negative for γ-H2AX foci (Fig. 2D and SI Appendix, Fig. S2E).

Treating cells carrying the mCherry/MYC-ER lentivirus with bleomycin indicated that any upregulation of reporter expression by DNA damaging agents other than topoisomerase inhibitors, was insufficient to achieve the high levels attained by non-damaging, low-dose CPT (SI Appendix, Fig. 2D). We surmise that the increased transcription activation and/or amplification elicited by low dose CPT primarily reflects altered DNA topology or conformation in response to topoisomerase inhibition.

If increased transcription amplification following TOP1 inhibition is not a consequence of DNA damage, but is due to interference with TOP1 activity, then other maneuvers that limit TOP1 action should yield similar results. Therefore, TOP1 or TOP2 were knocked down with siRNA, and cellular topoisomerase levels were verified by immunoblotting (Fig. 3A). mCherry expression was monitored in topoisomerase knockdown or control cells in the presence or absence of tamoxifen to activate MYC-ER. TOP1 knockdown dramatically increased mCherry fluorescence just as seen in cells treated with low-dose CPT (Fig. 3B). Knockdown of TOP1 increased the number and intensity of the mCherry-positive cells within both the zones of activation and amplification (Fig. 3B); amplification was dramatically enhanced by MYC-ER following treatment with tamoxifen (Fig. 3B). In contrast, the mCherry profile was much less affected by TOP2A knockdown (Fig. 3B). The increase in mCherry expression seen upon siTOP1 was confirmed at the RNA level by RT-qPCR (Fig. 3C). Thus, interfering with TOP1 activity, whether by low-dose CPT inhibition or siRNA knockdown, increased transcription amplification independent from DNA damage.

**Fig. 3.**
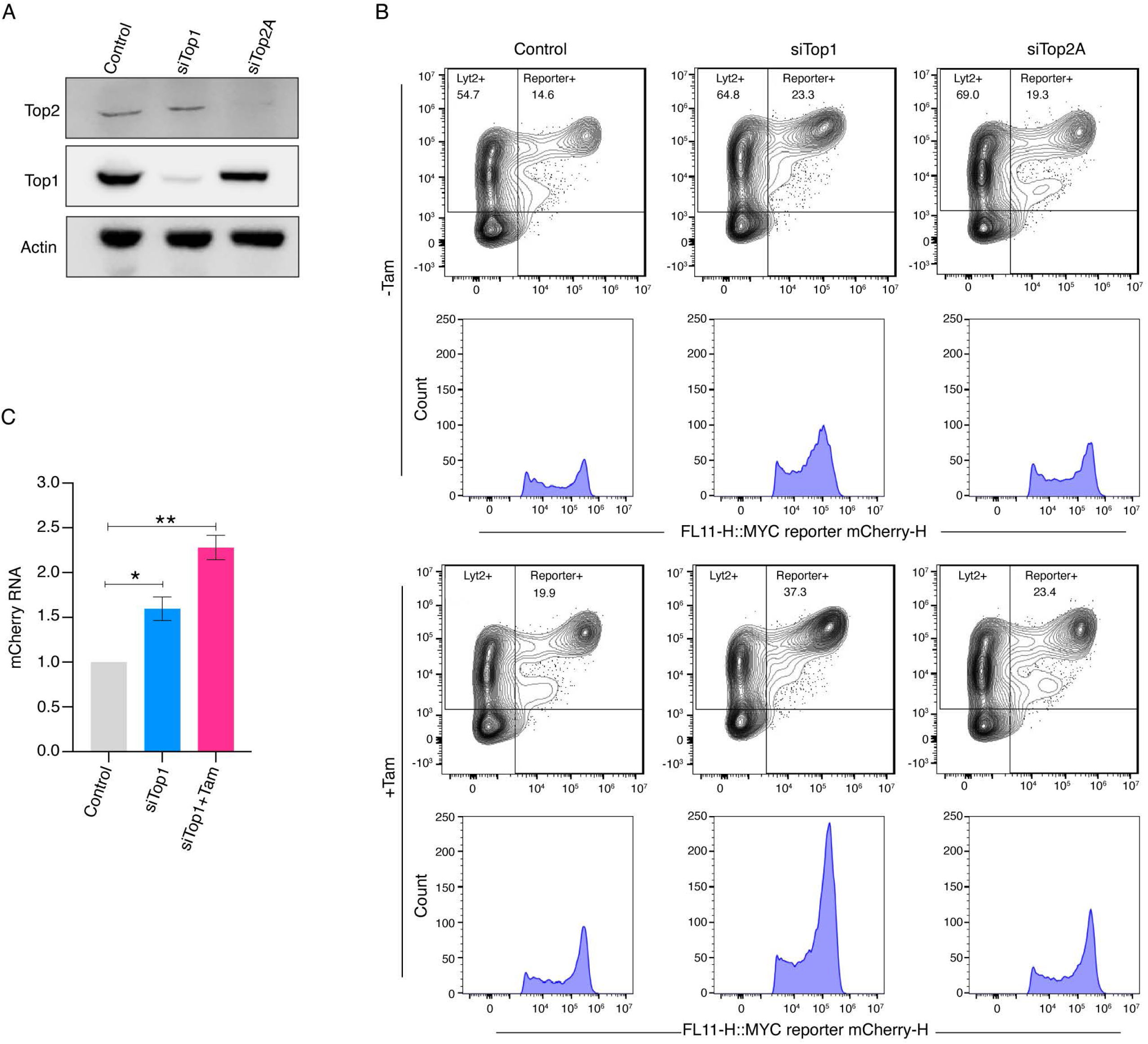
TOP1 knockdown augments transcription amplification by MYC. *(A)* Immunoblot showing TOP1 and TOP2 protein levels in cells treated with non-targeting or TOP1- and TOP2-specific siRNAs. Actin served as a loading control. *(B)* MCF10A cells expressing lentiviral E-box (+) mCherry constructs were treated with non-targeting or TOP1- and TOP2-specific siRNAs for 48 h, followed by Dox (150 ng/mL) and ± Tam (200 µM) treatment. Cells were analyzed by flow cytometry and mCherry fluorescence is shown on the X-axis, and LYT2 was detected using an anti-CD8α antibody on the Y-axis. Data were gated on live singlet cells gated for mCherry and LYT2 (Alexa-647) channels. Histograms below show fluorescence intensity distributions of mCherry-positive populations under each condition. *(C)* Quantification of fold change of mCherry mRNA expression in MCF10A cells with or without Tam (200 µM) upon siTOP1 treatment (calculated value ± SEM from three biological replicates). For all statistical analyses (t-test): *, and **, indicate P < 0.05, and 0.01, respectively.

### TOP1 inhibition potentiates MYC-dependent transcriptional amplification across the genome

Is the mCherry reporter’s response to topoisomerase inhibition generalizable across the genome? Transcriptional amplification by MYC is a global phenomenon augmenting the expression of all active genes. To determine whether the increased mCherry expression observed when TOP1 activity is inhibited or knocked down reflects a global effect or is instead specific to the lentivirus reporter system, RNA expression in these same cells was interrogated using RNA-seq.

Before exploring the genome-wide transcriptional response of MCF10A cells to treatment with low (100 nM) or high CPT (2 μM), we first confirmed by RTqPCR analysis that the increased mCherry expression in response to treatments with low dose CPT, and or tamoxifen occurred at the RNA level (Fig. 4A). Similarly, high-dose CPT interfered with Dox-activated mCherry expression at the RNA level (Fig. 4A) as expected (51). With every treatment, any upregulation by MYC-ER was prevented by inhibiting MYC-MAX dimerization (Fig. 4A). To interrogate the influence of CPT on global RNA expression, cells were treated with low or high concentrations of CPT for 4 or 24 hrs, both in the presence and absence of tamoxifen to activate MYC-ER, and differential gene was analyzed using DESeq2 (which ignores changes in total RNA levels between conditions). The analysis revealed a broad spectrum of responses; more genes were upregulated than downregulated, and some were left unchanged by drug treatment. No simple ontological patterns emerged here, it is important to reemphasize that DESeq2 reveals relative—not absolute--differential gene expression (7).

**Fig. 4.**
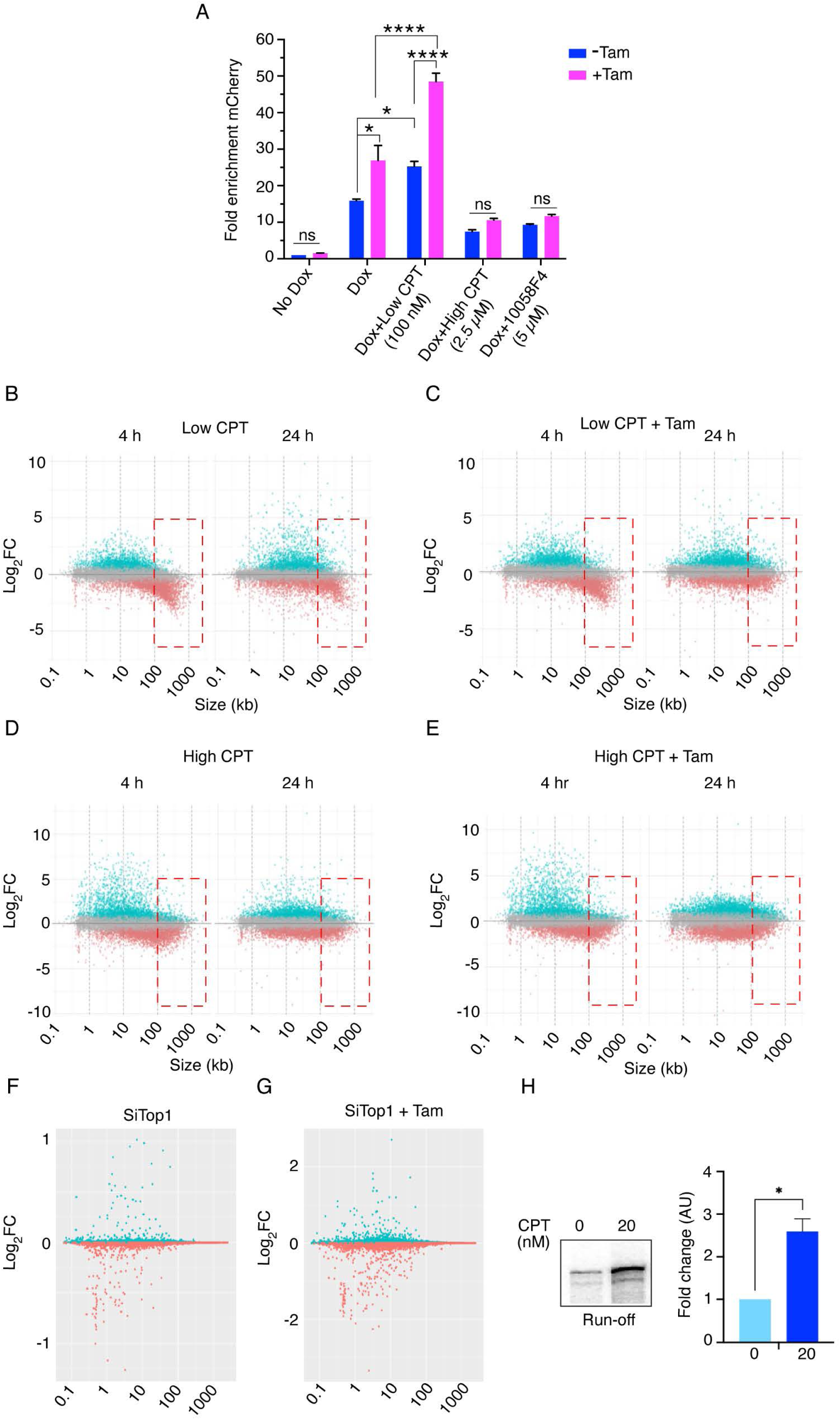
TOP1 inhibition potentiates MYC-dependent transcriptional amplification across the genome. *(A)* Quantification of fold change in mCherry mRNA expression in MCF10A cells expressing lentiviral E-box (–) mCherry constructs treated with various combinations of Dox (150 ng/mL), ± (200 µM), and either low (100 nM) or high (2.5 µM) camptothecin, or 10058-F4 (5 µM) (calculated as mean ± SEM from three biological replicates). For all statistical analyses (t-test): *, and ****, indicate P < 0.05, and 0.0001, respectively; ns denotes nonsignificant. *(B-E)* Log₂-fold change (Log₂-FC) versus gene length (kb) in RNA-seq data from MCF10A cells expressing lentiviral E-box (–) mCherry constructs. Cells were treated with various combinations of Dox (150 ng/mL), ± Tam (200 µM), and either low (100 nM) or high (2.5 µM) CPT. Plots show transcriptional responses at 4 h and 24 h post-treatment (in biological replicates), each point represents a single gene, with Log₂-FC derived from differential expression analysis (DESeq2) relative to untreated control. Gene length (kb) was plotted on the x-axis, and Log₂-FC on the y-axis. *(F-G)* Log₂-fold change (Log₂-FC) versus gene length (kb) in RNA-seq data from MCF10A cells expressing lentiviral E-box (–) mCherry constructs. Cells were treated with either non-targeting or TOP1-specific siRNAs for 24 h, followed by Dox (150 ng/mL), ± Tam (200 µM) for 24 h. Plots show transcriptional responses post-treatment (in biological replicates), each point represents a single gene, with Log₂-FC derived from differential expression analysis (DESeq2) relative to untreated control. Gene length (kb) was plotted on the x-axis, and Log₂-FC on the y-axis. *(H)* Run-off transcription from CMV-promoter–containing linear plasmid performed in the presence or absence of CPT (20 nM). The histogram represents the quantification of three independent experiments (mean ± SEM from three technical replicates). Asterisks indicate statistically significant differences (t-test, *P < 0.05).

Because gene length can influence transcriptional responses to topoisomerase inhibition, with prior studies showing relative over-expression of short genes and under-expression of long genes (41, 43, 44), we examined the effect of gene length on transcriptional responses to low- and high-dose CPT, as well as to TOP1 knockdown. Plotting the differential gene-expression against gene-length for all genes under the various conditions revealed that with low CPT treatment shorter genes appeared to be preferentially upregulated, whereas genes longer than ∼50 kb were more likely to be downregulated (Fig. 4B). Upon MYC activation, the upregulation of short genes by low CPT was exaggerated while the downregulation of long genes was attenuated, likely due to both MYC stimulation of the transcription machinery as well as MYC-enhanced topoisomerase 1 and 2 activities in the topoisome (Fig. 4C). The differential response to topoisomerase inhibition of short versus long genes, especially in the nervous system has been previously reported (41). A caveat to the interpretation reported in prior studies is that DESeq2 and other algorithms, routinely normalize RNA levels when comparing samples, so that the reported fold- up- or -downregulation is relative and not absolute. In other words, if topoisomerase inhibition globally depressed transcription, but depressed short gene expression less than it depressed long genes, then upon normalization, the short genes would score as being up-regulated despite not changing or even declining. However, the increase in the intensity of mCherry fluorescence seen in the low-CPT treated cells confirms an absolute as well as relative increase in the expression of this short of reporter. Perhaps the increased expression of short genes following treatment with low-level CPT was due to increased negative DNA supercoiling near promoters that facilitate promoter loading (30, 32, 35).

With high dose CPT, transcription was even more globally depressed as evidenced by an absolute decrease in mCherry fluorescence relative to untreated cells (Fig. 4A). At these higher doses, the preferential inhibition of long genes was less apparent (Fig. 4D). Most likely, at high levels of CPT, the inhibition of transcription was more uniform and independent of gene length (Fig. 4D). This transcriptional profile more closely resembled general inhibition of transcription induced by DNA damage (52, 53). Activation of MYC-ER by tamoxifen in cells treated with high CPT failed to coax higher expression from genes independent of their length (Fig. 4E) indicating that highly damaged DNA is poor substrate for transcription amplification by MYC (54). In contrast to mCherry, which was downregulated upon high CPT treatment (Fig. 4A, SI Appendix, Fig. S2B), genome-wide analyses revealed an apparent increase in the expression of short genes (Fig. 4D and 4E). This increase is likely due to normalization, as total RNA levels were reduced under high CPT conditions.

CPT intercalates at the interface between TOP1 protein and the cleaved DNA preventing ligation thus, stabilizing the covalent complex and its associated DNA nick. Within this complex, the rotation of the nicked strand around its partner is restricted, thus retarding the relaxation of supercoils (55–57). The release of positive supercoils is considerably more impaired than of negative supercoils as left-handed rotation of the double helix encounters more friction than right-handed. To ask whether the impaired transcription of long genes versus short genes was related to the impaired mechanics of the CPT-jammed TOP1 versus a general decrease in TOP1 activity (that would be compensated in considerable measure by the action of TOP2), mRNA expression was monitored following knockdown of TOP1 with siRNA. No differential effect of this knockdown on long genes versus short genes was noted (Fig. 4F and G). As in yeast, and in single molecule experiments, the poisoning of TOP1 with CPT returns a stronger phenotype than the reduction or elimination of TOP1B expression (57).

It is easy to understand why topoisomerase inhibition might interfere with gene expression as torsional stress builds up, and as knots and tangles accumulate in the template; but why would low-dose CPT increase the expression of some short genes? Because *in vitro* transcription becomes independent of TFIIE and TFIIH on negatively supercoiled templates, we reasoned that increased upstream negative supercoiling might facilitate promoter melting and PIC-formation, whereas topoisomerase activity later in the transcription cycle would promote pause-release and productive elongation (11, 15, 17, 58). Given the multiplicity of contacts of the DNA template with various components of the transcription machinery, associated transcription factors, and chromatin complexes, it is highly likely that during early stages of the transcription-cycle, torsional stress is partitioned between topological microdomains (59). If these microdomains generate or constrain torsionally-stressed segments of DNA that assist promoter melting and initiation, then relaxation due to premature topoisomerase activity might collapse the polymerase bubble and abort transcription. In this situation, topoisomerase inhibition might stabilize PICs and increase transcription. To test this notion *in vitro* transcription assays were conducted using a nuclear extract and a linear CMV-promoter driven template. These assays revealed that including 20 nM CPT in the reactions enhanced the run-off transcription product by 2-3-fold (Fig. 4H), similar to the stimulation seen *in vivo* (Fig. 4A). Note that this dose of CPT is less than that has been customarily tested in either *in vivo* or *in vitro* experiments. The results suggest that basal TOP1 activity near TSSs downregulates transcription.

### DNA supercoiling pre-amplifies MYC-mediated transcription at promoters

Psoralen intercalation assays were used to assess the topological response of the mCherry reporter following treatment with low concentrations of the TOP1 inhibitor CPT (60). Psoralen preferentially intercalates into negatively supercoiled DNA and, upon UV exposure, crosslinks the strands. qPCR was used to compare the level of psoralen cross-linking upstream of the mCherry promoter in the presence or absence of CPT, and as a function TetVP16 activation by doxycycline or switching on MYC-ER with tamoxifen (Fig. 5A). To confirm that any increase of psoralen incorporation reflected elevated negative DNA supercoiling, cells were also treated with bleomycin to introduce strand breaks that would help to relieve negative supercoiling thereby diminishing psoralen binding.

**Fig. 5.**
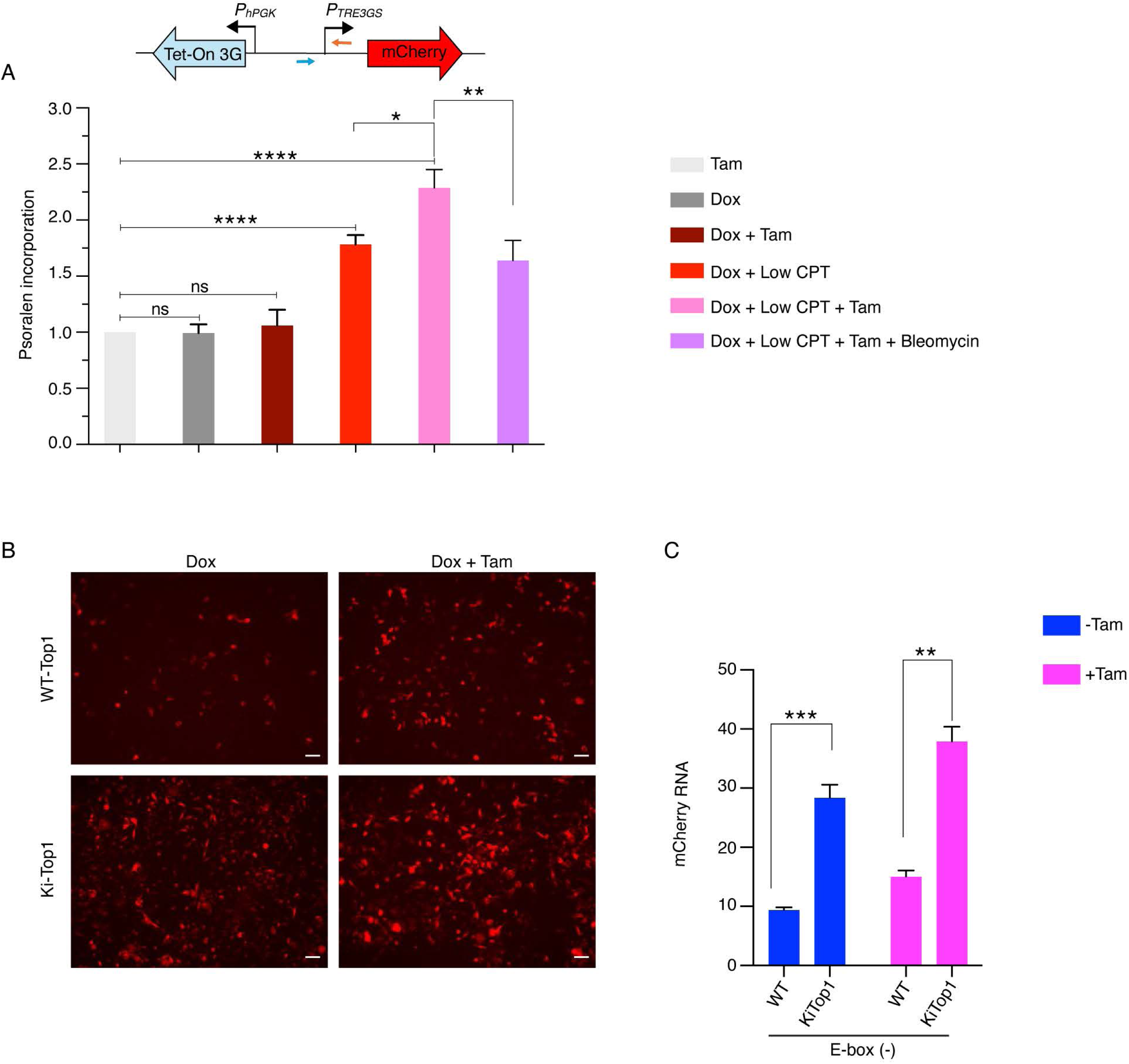
DNA supercoiling pre-amplifies MYC-mediated transcription at promoters. *(A)* Quantification of psoralen incorporation at the TRE3Gs (mCherry) promoter in MCF10A cells expressing lentiviral E-box (-) mCherry constructs following treatment with indicated combinations of Dox (150 ng/ml), Tam (200 µM), CPT (100 nM), and bleomycin (5 µM). Psoralen incorporation was quantified by qPCR, and data are presented as fold change relative to tamoxifen treated (calculated as mean ± SEM from six biological replicates). For all statistical analyses (t-test): *, **, and ****, indicate P < 0.05, 0.01,and 0.0001, respectively; ns denotes nonsignificant. *(B)* WT-TOP1 and KiTOP1 HCT116 cells expressing lentiviral E-box (-) mCherry constructs treated with the indicated conditions of Dox (150 ng/ml) and Tam (200 µM). Representative microscopic images from at least three independent experiments are shown (scale bars, 200 µm). *(C)* Quantification of fold change in mCherry mRNA expression in WT-TOP1 and KiTOP1 HCT116 cells expressing lentiviral E-box (-) mCherry constructs treated with the indicated conditions of Dox (150 ng/ml) and Tam (200 µM). Data represents mean ± SEM from three replicates. Asterisk denotes significant differences for, all statistical analyses (t-test): **, and ***, indicate P < 0.01,and 0.001, respectively.

Treatments with tamoxifen and/or Dox all supported similar constitutive levels of psoralen intercalation upstream of the mCherry minimal promoter (Fig. 5A). Note the torsional stress resident in this test segment would super-imposed dynamic negative supercoils propagating upstream from the oppositely oriented, powerful hPGK promoter on top of those emanating from the weaker mCherry promoter. Situated only ∼ 1kb upstream of the mCherry TSS, but oppositely oriented, the hPGK promoter maintains continuous expression of Tet-On 3G activator, and so generates constitutive dynamic negative supercoils that swamp out the lower level of torsion generated by the TREGS3 promoter (31, 32). Treatment with low levels of CPT augmented psoralen cross-linking in the test segment, and additional cross-linking accrued subsequent to MYC-ER activation with tamoxifen (Fig. 5A). These higher levels of psoralen binding were abolished by bleomycin treatment that releases torsional stress (Fig. 5A). These results suggest that the CPT-mediated build-up of promoter-localized negative supercoiling primes the formation of PICs that serve as substrates for whatever Dox- and TAM-enabled mechanisms enhance mCherry expression. If CPT increases mCherry promoter loading and expression, then TOP1 activity oppositely would be expected to dampen promoter melting and RNAPII reloading, while subsequently facilitating pause release and elongation (11).

To support the notion that interference with TOP1 activity increased RNAPII loading at promoters, we compared RNAPII levels at the mCherry promoter in HCT116 cells expressing wild-type or an exon-4–deleted variant of TOP1 (kiTOP1), which lacks the ability to interact with RNAPII (11). Disruption of this interaction prevents RNAPII-CTD from stimulating TOP1’s DNA- relaxation activity thereby leading to higher levels of negative DNA supercoiling and of RNAPII at promoters (61). HCT116kiTOP1 cells transduced with the MYC-ER-mCherry lentivirus were monitored for mCherry expression in the absence or presence of tamoxifen. The kiTOP1 cells supported greater mCherry expression than did cells with native TOP1, and expression was further increased upon MYC-ER activation with tamoxifen (Fig. 5B). These results support the notion that interference with TOP1 activity increases promoter loading, and so these loaded promoters provide more substrate for MYC’s actions. RT-qPCR confirmed that the increased mCherry fluorescence reflected increased mCherry mRNA (Fig. 5C). Notably, the results were similar irrespective of whether the mCherry promoter possessed an E-box (SI Appendix, Fig. 5A, 5B).

### Reduced TOP1 activity enhances RNAPII recruitment and facilitates transcription activation

We hypothesize that increased mCherry expression driven by low level CPT resulted from supercoil-enhanced promoter loading of RNAPII. In contrast, MYC-amplified transcription has generally been attributed to MYC-facilitated pause-release that redistributes RNAPII from promoters toward gene bodies. To examine the effects of CPT inhibition of TOP1 on RNAPII occupancy at the mCherry promoter, ChIP-seq and qPCR was performed. Low-dose CPT treatment increased the amount of RNAPII found at the mCherry promoter (Fig. 6A), suggesting inhibition of TOP1 enhanced promoter loading leading to upregulated expression (Note that if the RNAPII increase were attributed to ineffective pause release, then mCherry levels should have decreased—they increased). To ascertain whether low-dose CPT generally facilitated RNAPII loading at promoters, or whether this effect was somehow restricted to short genes, when the genome-wide profile of RNAPII at promoters was evaluated in response to low-dose CPT, the increase was found to be independent of gene length (Fig. 6B-D) as well as expression (SI Appendix, Fig. S6A-B). If promoter DNAs were more negatively supercoiled at steady-state when TOP1 was reduced, then RNAPII occupancy would be increased as promoter melting was facilitated. On short genes, this enrichment of RNAPII extended through their TESs, indicating successful full-length transcription despite CPT-interfering with removal of elongation-impeding positive supercoils (Fig. 6C, SI Appendix, Fig. S6C). In contrast, consistent with decreased expression of long genes upon low-CPT-treatment (Fig. 4B and 4C), RNAPII levels became progressively attenuated as the enzyme approached the 3’ ends of long genes (Fig. 6D, SI Appendix, Fig. S6D). So, the same treatment that elevated RNAPII levels at the TESs of short genes reduced RNAPII levels within TES regions of long genes. Although CPT interferes with the relaxation of both positive and negative supercoils, the effect on the former is far more profound, and so this effect is exacerbated on long genes (55, 57). That constitutive TOP1 activity is both antagonistic to promoter loading and conducive to elongation illustrates the importance of proper management of DNA mechanics and topology throughout the transcription cycle.

**Fig. 6.**
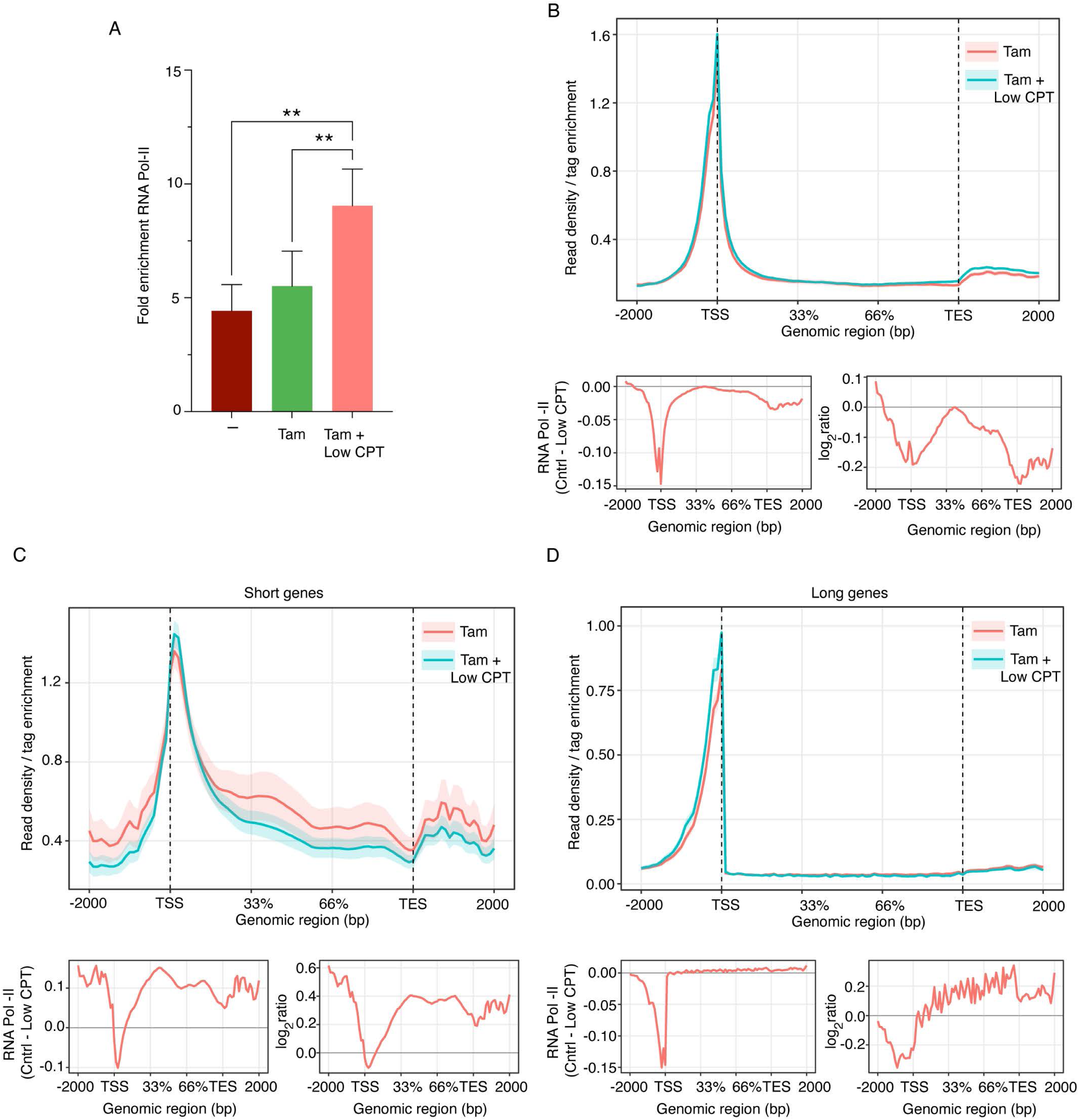
Reduced TOP1 activity enriches RNAPII at TSSs to facilitate transcription activation. *(A)* Fold-change in occupancy of RNAPII at the TRE3Gs (mCherry) promoter in MCF10A cells expressing lentiviral E-box (-) mCherry constructs following treatment with indicated combinations of Dox (150 ng/ml), Tam (200 µM) and CPT (100 nM) for 24 h. RNAPII occupancy was quantified by qPCR, and data are presented as fold change relative to IgG. Values represent mean ± SEM from three biological replicates. Asterisk denotes significant differences, for all statistical analyses (t-test): ** indicate P < 0.01. *(B-D)* Metagene profiles of RNAPII occupancy (read density per tag enrichment) in MCF10A cells expressing lentiviral E-box (-) mCherry constructs after 24 h treatment with indicated combinations of Dox (150 ng/ml), Tam (200 µM) and CPT (100 nM). Data are shown for all genes *(B)*, short genes (<25 kb) (C), and long genes (>50 kb) (D). The plot below each panel shows a metagene analysis of RNAPII occupancy depicting the difference between conditions (Control [Tam] – Low CPT [Tam + LowCPT] on the left, and the log₂ ratio of Control over Low CPT on the right. Signals are plotted across a normalized gene model spanning −2000 bp upstream of the transcription start site (TSS), through the gene body (33% and 66 % fractional lengths), and 2000 bp downstream of the transcription end site (TES). Positive values indicate higher RNAPII occupancy in Control condition, whereas negative values indicate enrichment in the Low CPT conditions.

## Discussion

Whether MYC is an activator or an amplifier of gene expression, transcription of its target genes requires chromatin decompaction, pre-initiation complex assembly, early transcription, pause release and elongation. RNAPII elongates along MYC responsive genes for distances short (as short as few hundred nucleotides--some pri-miRNA transcripts (62) and long, but with a median length of 24 kb. Likewise, loops of DNA-bound factors to promoter-bound RNAPII complexes, range from local promoter loops to promoter-enhancer interactions that span vast genomic distances. During active transcription, the DNA helix is at least transiently conformationally and topologically strained by promoter-melting, elongation, R-loop formation and resolution, and nucleosome gymnastics. All these processes introduce dynamic or static mechanical stresses into the double helix that may interfere with transcription or provoke DNA damage. DNA topoisomerases manage these stresses and resolve DNA knots and tangles that may arise in the template, even at the risk of topoisomerase-generated DNA damage (63). But whether topoisomerases are integral or auxiliary transcription-cycle participants is unknown. Topoisomerases may be essential for each round of the cycle, or they may be on standby to rectify topological and conformational issues as they arise. Similarly, MYC engages many partner proteins, including topoisomerases to increase gene expression (1, 8, 9, 54). But whether MYC participates in a stereotypical reaction pathway that includes topoisomerases to increase gene expression, or instead, if MYC acts opportunistically or stochastically to accelerate whatever step happens to be limiting at a given promoter, at a given time, is also unknown.

The experiments in this report shed light on MYC’s action throughout the transcription-cycle, on the participation of topoisomerases in basal transcription, and on MYC-activated transcription (Fig. 7). First, the lentivirus system herein cleanly distinguishes transcription activation from transcription amplification. Although doxycycline is strictly required to switch-on Tet-On 3G -driven mCherry expression, alone it does not achieve the same high levels of expression that are attained with active MYC (either endogenous or lentivirus -expressed). Similarly, tamoxifen alone fails to elicit any mCherry fluorescence, whether or not an E-box is included in the mCherry promoter to recruit MYC (endogenous or MYC-ER). Evidently, MYC cannot sponsor the same early steps of transcription enabled by Tet-On 3G. However, upon the treatment with Dox, the Tet-On 3G -activated mCherry promoter becomes a suitable substrate for MYC-driven transcription amplification. The efficiency of amplification by endogenous MYC and MYC-ER are notably enhanced by inclusion of a proximal promoter E-box demonstrating the functionality of this of element. Yet alone, this E-box is insufficient to activate transcription in the absence of another activator. We conclude that MYC operates later in the transcription-cycle than does Tet-On 3G.

**Fig. 7.**
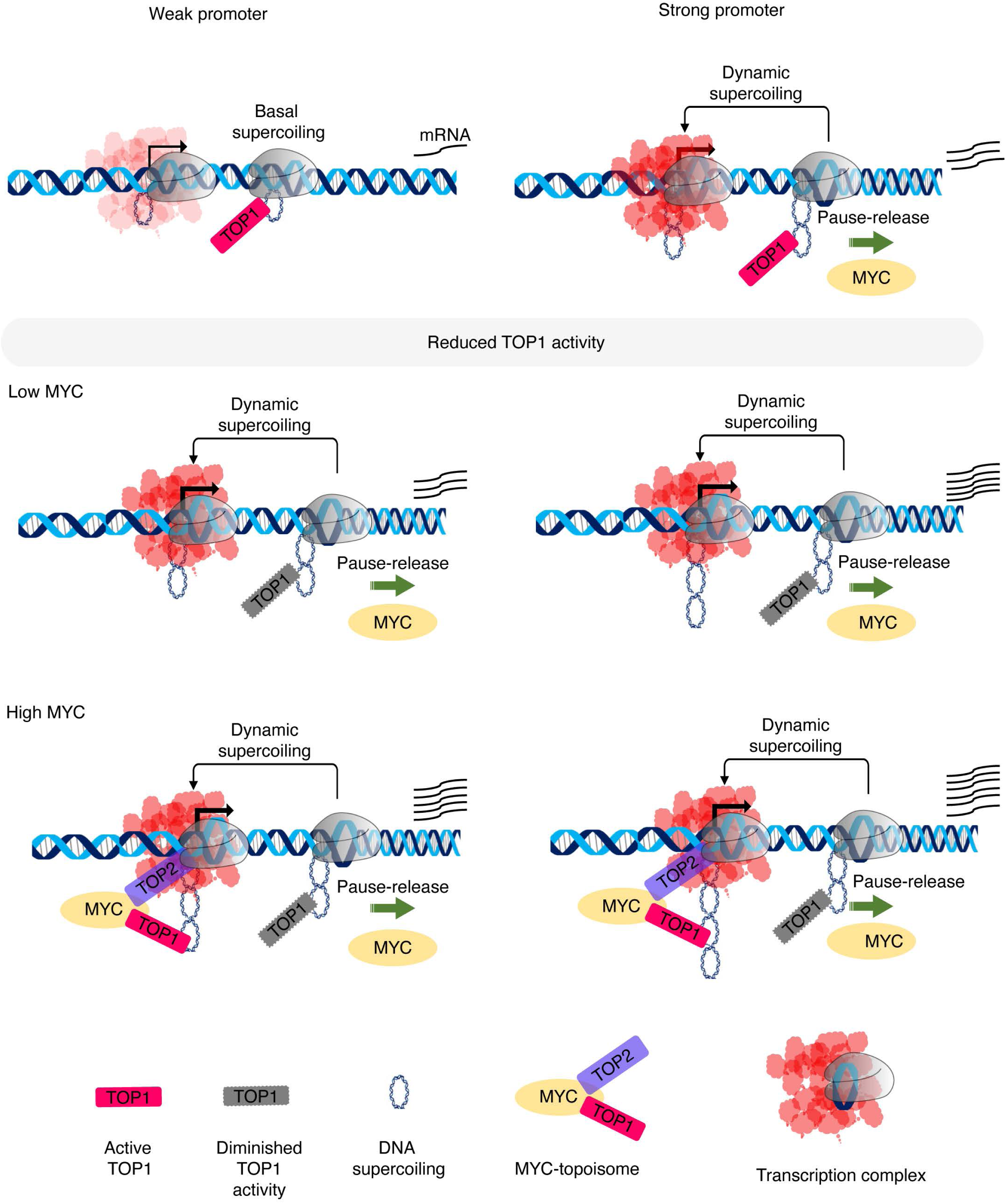
Reduced TOP1 activity facilitates promoter melting, PIC-formation, early transcription, and MYC-dependent transcription amplification. MYC preferentially amplifies highly-expressed genes maintaining high levels of dynamic supercoiling, while weak promoters lose static torsional stress through basal topoisomerase activity. Reduced TOP1 activity (CPT treatment or siTOP1) maintains negative torsional stress at transcription start sites (TSSs) facilitating promoter melting, RNAPII reloading, and early transcription, the substrate upon which MYC acts. In the absence of MYC, catalytically active TOP1 primarily engages downstream of pause sites; however, within the MYC-topoisome, TOP1 engagement shifts upstream to the TSS. By increasing baseline negative supercoiling, reduced TOP1 activity blurs the distinction between weak and strong promoters, limiting MYC’s selectivity.

Both the endogenous MYC-amplified and the MYC-ER amplified populations of mCherry positive cells are defined by and cohabit the same region of the mCherry/Lyt2-contour-plots. That tamoxifen treatment increased the number of cells in the amplified regions without further elevating the magnitude of mCherry fluorescence beyond the level supported by endogenous MYC, indicates the MYC driven output becomes saturated at an active promoter, and that in some cells, endogenous MYC levels are sufficient to saturate mCherry output. Thus, endogenous MYC-expression appears to be heterogeneous within the population of cells, but stable within individual cells as previously observed (64, 65). If short-lived MYC protein levels fluctuated rapidly within cells, saturation would not be maintained, and separation of the activated versus amplified populations would not be resolved.) As expected, if MYC is the major driver of amplification, both the endogenous MYC and MYC-ER amplified populations were sensitive to MYC inhibitors.

Low-dose CPT treatment alone does not activate mCherry expression but once activated by Dox, low-dose CPT treatment dramatically increases the amplified population in a manner resembling MYC; moreover, this increase is sensitive to MYC inhibitors. Thus, CPT creates a pre-amplified population that is primed to respond to MYC. We suggest that CPT retards the removal of negative supercoils at transcription start sites just as reported in single-molecule studies. As CPT-stabilizes the TOP1cc, it also increases resistance to rotation of the DNA within TOP1’s active site, thus retarding relief of torsional stress (55, 57). The resulting increase in unwinding stress at start sites would be expected to support promoter melting and RNAPII reloading thus providing more substrate for MYC to amplify. The CPT-provoked increased levels of RNAPII at promoters seen here, (both the mCherry promoter and genomic promoters-of both long and short genes) are in accord with this expectation.

Without MYC, free TOP1 becomes catalytically engaged downstream of pauses sites; but when included in the topoisome, MYC becomes immediately engaged at the TSS (11, 13). How can we rationalize this upstream shift of catalytically-engaged, topoisome-embedded TOP1? That CPT enhances transcription-amplification by endogenous MYC suggests an answer. In cells with low MYC, preservation of any resident TSS-localized supercoiling (arising from low level-transcription, chromatin mechanics, or DNA-conformational changes) would assist PIC formation and early transcription, and so the stimulation of TOP1-activity post-pause release facilitates both initiation and elongation. At weakly expressed promoters, torsional stress entropically decays or is released by basal topoisomerase activity before it can be harvested for reinitiation whereas during bursts at more highly expressed genes, promoters become dynamically strained facilitating melting and reinitiation. MYC has been proposed to operate preferentially on highly expressed genes until their output saturates (5–7, 66).

By engaging topoisomerases at TSSs, MYC helps to remove baseline, static supercoiling at both weak and strong promoters, but because dynamic supercoiling is relatively refractory to removal by topoisomerases (31, 32, 67), topoisome engagement at the TSS, attenuates transcription of the former, while dynamic supercoiling helps to sustain bursting and reinitiation of the latter (31, 32, 67). By increasing the baseline level of negative supercoiling due to topoisomerase inhibition, CPT blurs the distinction between lowly and highly expressed promoters (5–7, 66). Thus, CPT lessens the ability of MYC to discriminate high-output from weak-output promoters. In the absence of the drug, the removal of static torsional stress from TSSs impairs promoter reloading and MYC recruitment at weak promoters, while highly expressed genes that generate dynamic supercoils support a milieu conducive for MYC action. Hence, the upstream shift of TOP1 activity within the topoisome would sharpen the distinction between low expressed genes and bona fide MYC targets expanding the dynamic range of transcription amplification by MYC (Fig. 7).

If RNAPII promoter loading is increased at both short and long genes following TOP1 inhibition by CPT, then why does the output of the former increase while that of the latter decrease? The mechanics of TOP1 explain this difference. Single molecule studies have established that CPT impairs the relaxation of positive more than negative supercoils, and that it reduces transcription elongation rates by as much as two-thirds. Even with this slowed elongation, the RNAPII is able to complete the transcription of a short gene, whereas a long gene is more likely to develop RNA loops, DNA-damage, and other encumbrances that prevent completion of the primary transcript (68). Indeed, we find an increase in RNAPII at the TESs of short genes following CPT treatment (likely reflecting the increase in promoter loading), whereas fewer RNAPIIs reach the TESs of long genes (likely due to increased positive supercoiling and the consequences thereof).

Study of the interplay between DNA topology, dynamic supercoiling, RNAPII mechanics and MYC-action may reveal new insights into physical constraints on transcription and expose new vulnerabilities in cancer. The system described herein promises to provide a platform to interrogate genetically and pharmacologically the process of transcription amplification by MYC; the results of these explorations promise to increase our understanding of the physiological and pathological roles of MYC.

## Materials and methods

### Cell culture and drug treatment

MCF10A human mammary epithelial (ATCC, CRL-10317), HCT116 Human colon carcinoma (ATCC, CCL-247), HCT116KI (11) and OCI-LY1 (RRID: CVCL_1879) cell lines were used in this study. MCF10A cells were cultured in phenol red free DMEM-F12 (Invitrogen, 11-039-047) using media formulations prepared with or without EGF and serum, depending on the experimental condition. The complete growth medium consisted with DMEM/F12 supplemented with 2.5% horse serum (R&D Systems, S12150), 20 ng/mL EGF (Peprotech, AF-100-15), 10 µg/mL insulin (Sigma-Aldrich, I1882-100MG), 0.5 mg/mL hydrocortisone (Sigma-Aldrich, H0888-1MG), 100 ng/ mL cholera toxin (Sigma-Aldrich, C8052-1MG), 100 U/mL penicillin- streptomycin (Gibco, 15-140-163). MCF10A cells were cultured in complete growth medium until reaching approximately 60–70% confluence. Cells were then washed and incubated in complete medium lacking serum and EGF for 24 h prior to treatment with the indicated concentrations of Dox and Tam. HCT116 and HCT116KI cells were cultured in high glucose (4.5g/L) DMEM (Thermo Fisher, 11965092), supplemented with MEM nonessential amino acids (Corning, 25-025-Cl), 100 U/mL penicillin- streptomycin (Gibco, 15-140-163), and 10% bovine serum (Thermo Fisher, 26170043). OCI-LY1 cells were maintained in advanced RPMI 1640 (Gibco 12633-012) or RPMI 1640 (Gibco #61870-036) supplemented with 5% bovine serum, 100 U/mL penicillin- streptomycin (Gibco, 15-140-163) and 1% L-glutamine. HEK293T and HEK293FT cells were cultured in DMEM supplemented with 10% 10% bovine serum (Thermo Fisher, 26170043) and 100 U/mL penicillin- streptomycin (Gibco, 15-140-163) and 1% L-glutamine. All cell lines were maintained in a humidified incubator with 5% CO_2_ at 37°C and routinely tested and confirmed to be free of mycoplasma contamination.

### Plasmid construction

The pLVX-TetOne-Puro Vector (ClonTech, #631849) was used as the backbone for generating constructs expressing MYC-ER, LYT2 and mCherry. The mCherry coding sequence was amplified from the plasmid pLVX-PKG-GAL4USA-(Ebox)-E1bTATA-mcherry-hygro (5) using sequence-specific primers and inserted downstream of TRE3Gs promoter at *EcoR1* and *BamH1* restriction sites. To generate a promoter variant containing an E-box regulatory element, an E-box motif (CACGTG) was introduced into the TRE3GS promoter to create the +E-box version.

The original puromycin resistance cassette in plasmid pLVX-TetOne-Puro was replaced with LYT2 (mouse CD8a) driven by the SV40 promoter. A sNRP1polyA sequence was cloned downstream of the WPRE element to stabilize transcription termination.

The EF1-α promoter sequence was amplified from pKLV-EF1a-mCherry-W (Addgene #159295) and inserted into the vector at the pLVX-TetOne *KpnI* and *AgeI* sites. The MYC-ER fusion sequence was amplified from pRRL.Sin.cPPT.SFFV-MYC-ER_IRES-Tomato.WPRE (a gift from Martin Eilers) and inserted under the control of EF1- α promoter. The final constructs were designated pLVX-TetOne-Lyt2-Mcherry-EF1A_MYC-ER_NoE-box and pLVX-TetOne-Lyt2-Mcherry-EF1A_MYC-ER_+Ebox. Primers used for cloning are listed in Table S1.

### Cell line manipulation and generation

Lentiviral packaging plasmids pMD2.G (Addgene, #12259), psPAX2 (Addgene, #12260), together with the expression constructs pLVX-TetOne-Lyt2-Mcherry-EF1A_MYC-ER_NoE-box or pLVX-TetOne-Lyt2-Mcherry-EF1A_MYC-ER_+Ebox, were used to generate stable cell lines. Lentivirus particles were produced in HEK293FT cells using the TransIT293 transfection reagent and Opti-MEM medium, virus containing supernatants were collected and used to infect MCF10A, HCT116 and OCI-LY1 cells. The transduced cells were sorted by flow cytometry by staining cells for positive LYT2 surface expression (Alexa Fluor 647 -conjugated anti-mouse CD8a antibody) and mCherry fluorescence.

### FACS staining and analysis

Cells were treated with indicated drugs, including doxycycline, tamoxifen, CPT, etoposide, ICRF-187. Following treatment for the specified durations, cells were washed once with FACS buffer (2%FBS and 1mM EDTA in PBS) and stained with Alexa Fluor 647 - conjugated anti-mouse CD8a antibody (1:1000 dilution) for 20 min at 4 °C in the dark. After staining, cells were washed twice with FACS buffer to remove unbound antibody. Stained cells were analyzed on a Beckman Coulter CytoFlEX S flow cytometer to quantify surface LYT2 expression and mCherry fluorescence. Data were acquired using CytoFlex software and analyzed with FlowJo.

### siRNA transfection

All siRNA transfections were performed according to the manufacturer’s instructions using the DharmaFECT Transfection Reagent (Horizon Discovery). In brief, cells were seeded in 6 well plates 24 hr before siRNA transfection. Before transfection, growth media was changed, and cells were washed with PBS. 20 nM of each siRNA TOP1 siRNA (DharmaconTM, M-005278000005), and TOP2A siRNA (DharmaconTM, M-004239020005) (DharmaFECT) along with non-targeting control siRNA (DharmaFECT) was mixed with 4 ul of DharmaFECT®-2 reagent and incubated at room temperature for 10 minutes. siRNA mixes were then added to the cells having 2 ml of Opti-MEM antibiotics-free medium and incubated at 37°C, 5% CO2. After 8 hrs of incubation, Opti-MEM media was replaced with normal growth media. After 48hr post-siRNA transfection, cells were treated with indicated drugs for specific time.

### Quantitative Real Time PCR

Total RNA was isolated from 2 × 10⁶ cells using the Direct-zol™ RNA Miniprep Kit (Zymo Research, R2051-A) according to the manufacturer’s instructions. All samples were treated with the supplied DNase I during the purification to remove genomic DNA. For cDNA synthesis, 1µg of total RNA was reverse-transcribed using the SuperScript™ III First-Strand Synthesis System (Invitrogen, 18080-051) along with the Oligo DT (Invitrogen, 55063) and random hexamer (Invitrogen, 100026684), following the manufacturer’s protocol. Quantitative real-time PCR was performed using iTaq Universal SYBR Green Supermix (Bio-Rad, 1725121) on a LightCycler® 480 Instrument II (Roche). Each 12-µl reaction contained 6 µl of 2× SYBR Green Supermix, 1 µl of 10-fold diluted cDNA, and primers at a final concentration of 0.5 µM. Thermocycling conditions were as follows: initial denaturation at 50 °C for 2 min and 95 °C for 5 min, followed by 45 cycles of 95 °C for 10 s, 60 °C for 1 min, and 72 °C for 30 s. A melt-curve analysis was performed at 95 °C for 10 s and 60 °C for 1 min. Relative gene expression was determined using β-actin as the reference gene, with values were normalized to those of untreated cells. Primer sequences are provided in Table S2.

### Western blotting

Cells were washed twice with PBS, scraped and lysed in RIPA buffer (10 mM Tris-HCl pH 8, 150 mM NaCl, 0.5 % Sodium deoxycholate, 0.1 % SDS, 1 % NP40) supplemented with complete protease inhibitors. Lysates were vortexed and incubated for 1hr at 4°C, followed by centrifugation at 14000 rpm for 10 min at 4°C. The supernatants were collected and stored at −80°C. Equal amount of total protein were resolved on 4-12% Bis-Tris NuPAGE gels (Thermo Fisher, NP0323), transferred to nitrocellulose membranes, and probed with the primary antibodies as indicated. Proteins were detected by chemiluminescence using ECL (Thermo Scientific) and imagined with an Odyssey infrared scanner (Li-Cor). The reagents and sources are listed in Table S3.

### Immunofluorescence Staining and Confocal Microscopy

Immunofluorescence staining and confocal imaging were performed as previously described (12). Briefly, 1×10^4^ cells were seeded onto Ibidi 15 μ-slide angiogenesis chambers (ibidi-treat, 81506) and treated with the indicated drugs, followed by fixation in 4% paraformaldehyde for 20 min at room temperature. Cells were stained with γH2AX primary antibody (1:200), and Alexa Fluor 488/568-conjugated anti-mouse IgG secondary antibodies (1:1000; Invitrogen). Nuclei were counterstained with DAPI (Vector Laboratories) and images were acquired on a Zeiss LSM880 confocal microscope equipped with a 63×/1.4 NA oil objective. Images were processed in ImageJ (Fiji) and sized in Adobe Photoshop 7.0. γH2AX intensity per nucleus was quantified as total fluorescence normalized to cell number. Data visualization and statistical analyses were performed using GraphPad Prism 9.1.0. with two-way ANOVA and Dunnett’s correction for multiple comparisons. Significance is indicated as follows: p < 0.05 (*), p < 0.01 (**), p < 0.001 (***), and p < 0.0001 (****).

### *In vitro* transcription assays

*In vitro* transcription assays were performed as described previously (69, 70). Preinitiation complexes (PICs) were assembled by incubating 200 ng of linearized CMV promoter template (pGL2-CMV), 40 mg Hela nuclear extract, 10 mL H_2_O, and 7.5 mL HM.1 (20 mM HEPES, pH 7.5, 100 mM KCl, 12.5 mM MgCl_2_, 20 % glycerol), without or with CPT (20 nM), for 30 min at room temperature. Pulse transcription was initiated by addition of 0.5 mL [α-^32^P]CTP (3000 Ci/mmol) and 0.5 mM rGAC for 30 s, followed by a chase with 3 mM CTP for 5 min. Reactions were terminated by addition of stop buffer (100 mM Tris pH 7.5, 100 mM NaCl, 10 mM EDTA, 1% sarkosyl) and extracted with an equal volume of phenol/chloroform/isoamyl alcohol. The aqueous phase was precipitated with 1/10 volume of 5 M ammonium acetate, glycogen, and ethanol at −80 °C for 15 min, followed by centrifugation at 15,000 rpm at 4 °C. RNA pellets were resuspended in RNA loading buffer, heated at 75 °C, cooled on ice, and resolved on 8 % denaturing urea/TBE polyacrylamide gels. Dried gels were exposed to phosphorimager screens and analyzed using a Cytiva/Amersham Typhoon phosphorimager.

### RNA-sequencing

MCF10A-MYC-ER_NoBox cells were treated in various conditions of drugs or siRNA as indicated for various time points. Total RNA was extracted from 2×10^6^ cells using the Direct-ZolTM RNA Miniprep kit (15596018, Zymo Research, R2051-A), as described above. Libraries were prepared using the Illumina Total RNA Ligation Kit with RiboZero Plus according to the manufacturer’s instructions. Briefly, ribosomal RNA (rRNA) was depleted using biotinylated, target-specific oligonucleotides in combination with Ribo-Zero rRNA removal beads. The resulting rRNA-depleted RNA was fragmented into short fragments and reverse-transcribed into first-strand cDNA using random primers and reverse transcriptase. Second strand cDNA synthesis was subsequently carried out using DNA Polymerase I and RNase H. The resulting double-strand cDNA was used as input for a standard Illumina library preparation, including end-repair, adapter ligation, and PCR amplification to generate sequencing ready libraries. Final libraries were purified and quantified by qPCR prior to cluster generation and sequencing on NovaSeq 6000 platform.

The data was analyzed as follows, GEO accession number (). Adapters were trimmed (Cutadapt). Reads were aligned to hg38_lentivirus with STAR (avg. 92% uniquely mapped; 58–76% non-duplicates). Gene-level counts were produced with feature Counts; genes with < 80 total reads across the study were removed. Analysis was done using DESeq2. All code for RNA-seq analysis is available on GitHub [https://github.com/pop3www/GMS_Ranking]

### Chromatin Immunoprecipitation (ChIP) sequencing

The binding of RNAPII at the TRE3Gs (mCherry) promoter was performed with the ChIP-IT High Sensitivity kit (Active Motif, 53040) following the manufacturer’s instructions with minor modifications. Briefly, 2×10^7^ cells were fixed in the growth medium containing 1% formaldehyde and fixation was quenched with 0.125M glycine. Cells were chilled and sonicated using a Diagenode Bioruptor (low amplitude, pulse 30-s on/30-s off cycle for a total on time of 10–20 min) to obtain chromatin fragments averaging 200-600 bp. Chromatin immunoprecipitation was performed using 2 µg of anti-RNAP II antibody (Active motif, 102660) or normal rabbit IgG (Santa Cruz Biotechnology, sc-2017) as a negative control. Antibody–chromatin complexes were captured using 40 µl of Protein A/G magnetic beads (Pierce, 88803) and incubated at 4 °C for 4 h with gentle rotation. ChIP-DNA was then purified using the provided ChIP Filtration Columns as per manufactured instructions.

Libraries were prepared using the SWIFT_Accel-NGS 2S plus DNA Library Prep kit according to the manufacture’s protocol. Briefly, the 5′ and 3′ ends of ChIP DNA were dephosphorylated to prevent chimera formation and enhance adapter ligation efficiency to 3′ ends. Additional 3′ end repair was performed, along with polishing of both 3′ and 5′ overhangs. The i7 indexed adapter were ligated to the 3′ ends followed by 5′ end repair and ligation of the i5 indexed adapter to 5′ ends of the dsDNA substrate. SPRI bead-based purification steps were used to remove excess oligonucleotides and short fragments, as well as to exchange enzymatic buffer between steps. The adaptor-ligated DNA was PCR-amplified to enrich fragments containing adapters on both ends. Final libraries were purified, quantitated by qPCR, and followed by cluster generation and sequencing on the Illumina NextSeq platform. The data was analyzed as follows, GEO accession number (). All code for ChIP-seq analysis is available on GitHub [https://github.com/pop3www/GMS_Ranking].

### Quantitative PCR (qPCR)

qPCR was carried out using iTaq Universal SYBR Green Supermix (Bio-Rad, 1725121) on a LightCycler 480 Instrument II (Roche). Each 12-µl reaction contained 6 µl of 2× SYBR Green master mix, 0.5 µM of each primer, and 2 µl of ChIP or input DNA.

Thermocycling conditions consisted of initial denaturation (50°C for 2 min and 95°C for 5 min), followed by 45 cycles of amplification (95°C for 10 sec, 60°C for 1 min, and 72°C for 30 sec) and a melting step (95°C for 10 sec, 60°C for 1 min). The relative occupancy of RNAP II at TRE3Gs (mCherry) promoter was quantified by determining its fold enrichment over the IgG control. Primer sequences are provided in Table S2.

### Psoralen photobinding and qPCR

Psoralen photobinding to determine the DNA supercoiling in cells was carried out with modification from a previously described method (30, 71). In brief, cells (2×10^6^ in 6 mm dishes) were treated with 6 ug/ml of 4,5′,8-trimethylpsoralen (MP Biomedicals) for 5 min at 37 °C in the dark. Plates with cells were exposed to 3.6 kJ m−2 of 365-nm light (UV lamp, model B-100 A, Ultra-Violet Products). Cross-linked genomic DNA was isolated by RNase and proteinase-K treatment in lysis buffer (10 mM Tris-Cl, pH 8.0, 100 mM EDTA, 0.5% SDS), followed by phenol-chloroform extraction and ethanol precipitation. Purified DNA was sonicated (Ultrasonic processor XL sonicator, MISONIX, at 15% of power) to produce 250-bp-average-size DNA fragments. Cross-linked fragments were enriched by two rounds of denaturation and exonuclease I (Exo I, NEB) digestion. The remaining crosslinked duplexes were further purified by single round of denaturation and exonuclease VII (Exo VII, NEB) digestion. After final DNA purification, its concentration was assayed by Qubit fluorimeter (Invitrogen). The specificity of the crosslinked DNA purification was evident upon negligible recovery of DNA upon parallel treatment of DNA from the cells not exposed to UV light. Enrichment of specific DNA loci was determined by qPCR as described above with primers specific to TRE3Gs promoter. The reagents and sources are listed in Table S3.

## Acknowledgments

This research was supported by the Intramural Research Program of the National Institutes of Health (NIH). The contributions of the NIH authors are considered works of the United States Government. The findings and conclusions presented in this paper are those of the author(s) and do not necessarily reflect the views of the NIH or the U.S. Department of Health and Human Services. RNA-seq and ChIP-seq libraries were prepared and sequenced by the NCI Sequencing Facility and Genomics Core.

## Author contributions

Conceptualization, D.L, R.K.J.; Methodology, most of the experiments was performed by R.K.J., F.K., J.D.P., S.K.D.; Computational Analysis, B.W.; Investigation, D.L., R.K.J., F.K., J.D.P., S.K.D.,B.W. B.A.L; Resources, D.L; Writing – Original draft, R.K.J., D.L.; Writing – Review and editing, D.L., R.K.J., F.K., J.D.P., S.K.D.,B.W. B.A.L.; Visualization, R.K.J., F.K., B.W., S.K.D.; Supervision, D.L.; Funding Acquisition, D.L.

## Supporting Information

**Fig. S1.**
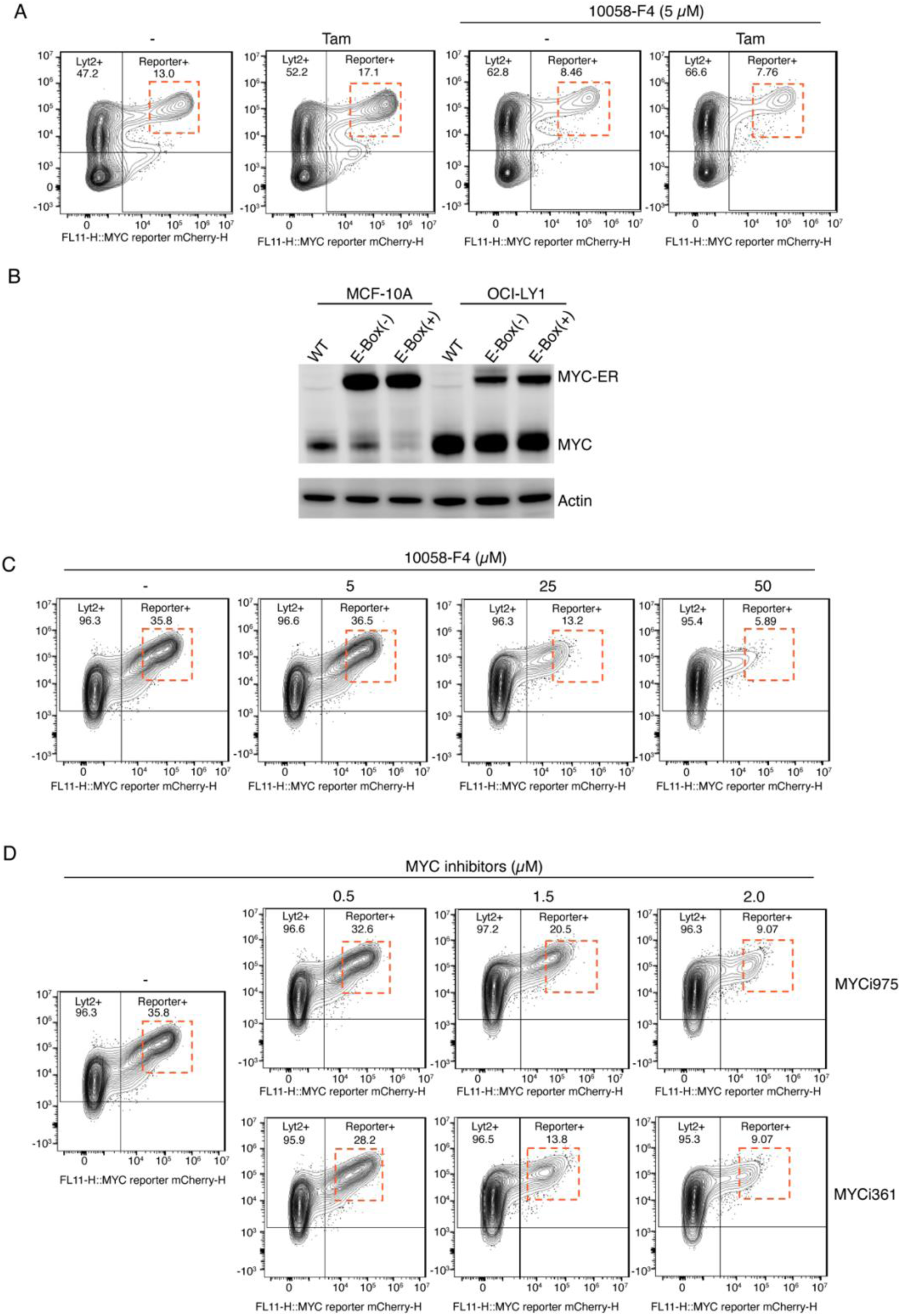
A cell-based assay establishes MYC as a transcription amplifier. *(A)* MCF10A cells expressing lentiviral E-box+ mCherry constructs were cultured under indicated Dox/Tam conditions ± MYC inhibitor 10058-F4 (5 µM). Cells were analyzed by flow cytometry and mCherry fluorescence is shown on the X-axis, and LYT2 was detected using an anti-CD8α antibody on the Y-axis. Data were gated on live singlet cells gated for mCherry and LYT2 (Alexa-647) channels. Transcriptionally amplified populations are highlighted with red dashed box. *(B)* Immunoblot showing MYC and MYC-ER protein levels in MCF10A and OCI-LY1 cells treated with Dox (150 ng/ml) and Tam (200 µM). Actin served as a loading control. *(C)* OCI-LY1 lymphoma cells expressing lentiviral E-box+ mCherry constructs were under indicated Dox/Tam conditions ± MYC inhibitor 10058-F4 (5 −50 µM). Cells were analyzed by flow cytometry and mCherry fluorescence is shown on the X-axis, and LYT2 was detected using an anti-CD8α antibody on the Y-axis. Data were gated on live singlet cells gated for mCherry and LYT2 (Alexa-647) channels. Transcriptionally amplified populations are highlighted with red dashed box. *(D)* OCI-LY1 lymphoma cells expressing lentiviral E-box+ mCherry constructs were cultured under indicated Dox/Tam conditions ± MYC inhibitors MYCi975 or MYCi361 at various concentration (0.5 −2 µM). Cells were analyzed by flow cytometry and mCherry fluorescence is shown on the X-axis, and LYT2 was detected using an anti-CD8α antibody on the Y-axis. Data were gated on live singlet cells gated for mCherry and LYT2 (Alexa-647) channels. Transcriptionally amplified populations are highlighted with red dashed box.

**Fig. S2.**
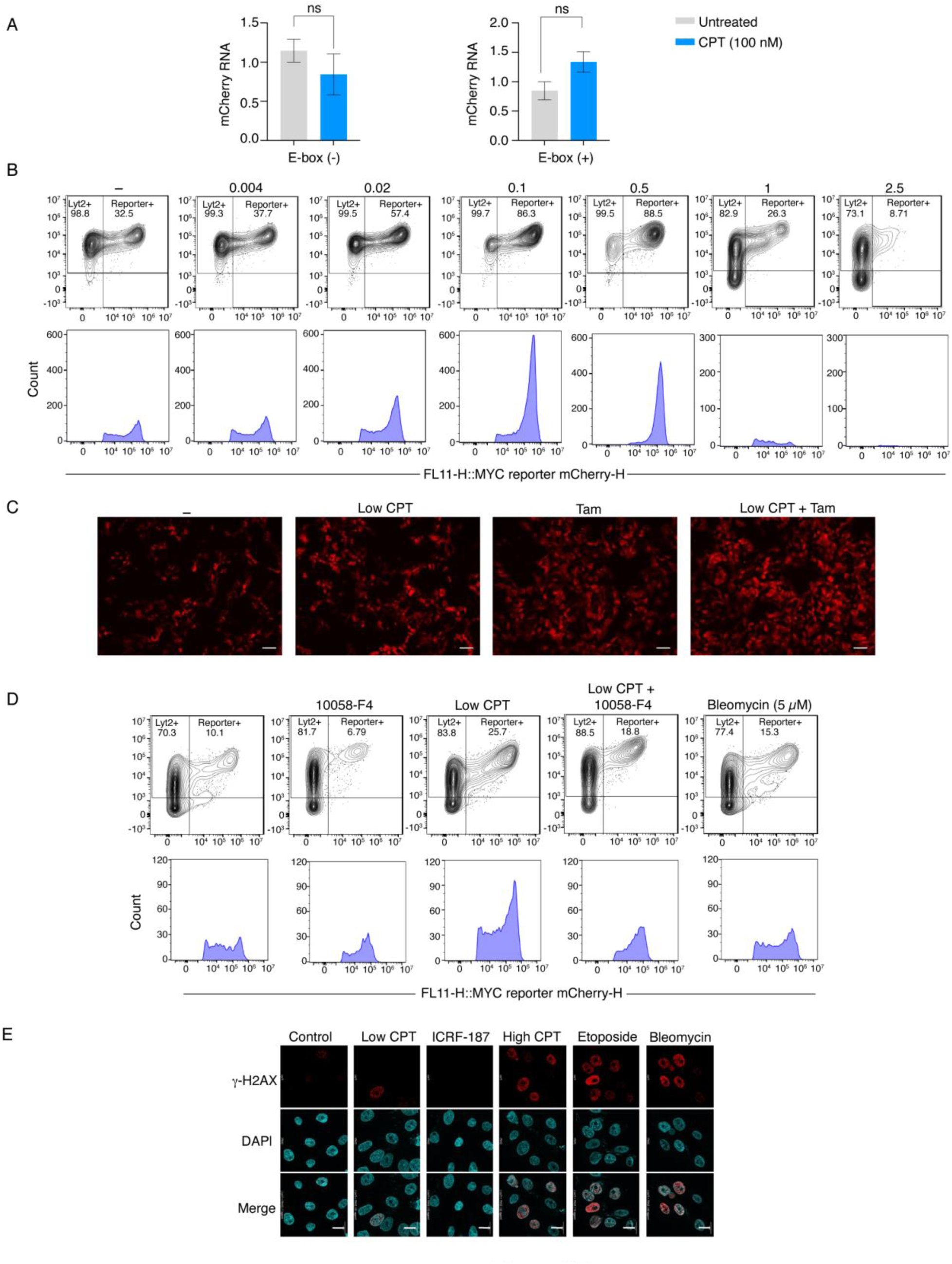
Inhibition of TOP1 enhances transcription amplification by MYC, independent of DNA damage. *(A)* MCF10A cells expressing lentiviral mCherry constructs with or without E-box were treated with camptothecin (100 nM) and mCherry expression was quantified by RT-q-PCR (t-test, calculated values ± SEM from three biological replicates). *(B)* MCF10A cells expressing lentiviral E-box+ mCherry constructs were treated with Dox (150 ng/mL) ± Tam (200 µM) and treated with the indicated concentration of camptothecin. Cells were analyzed by flow cytometry and mCherry fluorescence is shown on the X-axis, and LYT2 was detected using an anti-CD8α antibody on the Y-axis. Data were gated on live singlet cells gated for mCherry and LYT2 (Alexa-647) channels. Histograms below show the fluorescence intensity distributions of mCherry-positive populations under each condition. *(C)* MCF10A cells expressing lentiviral E-box (–) mCherry constructs were cultured in the presence of Dox (150 ng/ml) in the presence or absence of Tam (200 µM) and CPT (100 nM). Representative microscopic images from at least three independent experiments are shown (scale bars, 200 µm). *(D)* MCF10A cells expressing lentiviral E-box (+) mCherry constructs were cultured with Dox (150 ng/mL) and treated with indicated combination of CPT (100 nM), 10058-F4 (5 µM), and bleomycin (5 µM). Cells were analyzed by flow cytometry and mCherry fluorescence is shown on the X-axis, and LYT2 was detected using an anti-CD8α antibody on the Y-axis. Data were gated on live singlet cells gated for mCherry and LYT2 (Alexa-647) channels. Histograms below show the fluorescence intensity distributions of mCherry-positive populations under each condition. *(E)* Immunofluorescence microscopy of MCF10A cells expressing lentiviral E-box (+) mCherry constructs under treatment with Dox (150 ng/mL) ± Tam (200 µM) and treated with the indicated topoisomerase inhibitors- low (100 nM) versus high dose CPT (2.5 µM), etoposide (20 μM), ICRF-187 (5 μM), and bleomycin (5 μM). γH2AX foci are red. Nuclei were stained with DAPI (scale bars, 25 μm).

**Fig. S5.**
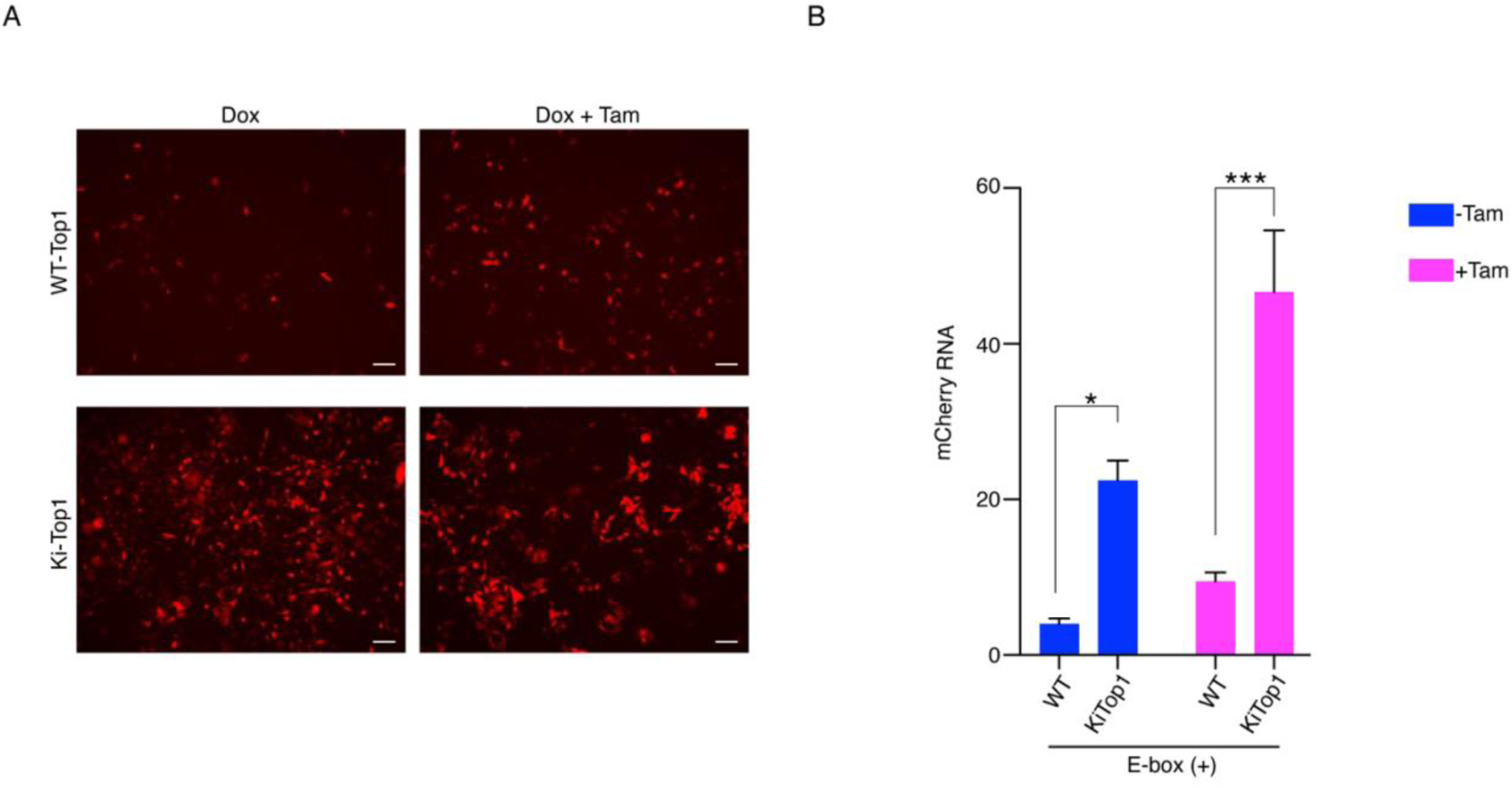
DNA supercoiling pre-amplifies MYC-mediated transcription at promoters. *(A)* WT-TOP1 and KiTOP1 HCT116 cells expressing lentiviral E-box (+) mCherry constructs treated with the indicated conditions of Dox (150 ng/ml) and Tam (200 µM). Representative microscopic images from at least three independent experiments are shown (scale bars, 200 µm). *(B)* Quantification of fold change in mCherry mRNA expression in WT-TOP1 and KiTOP1 HCT116 cells expressing lentiviral E-box (+) mCherry constructs treated with the indicated conditions of Dox (150 ng/ml) and Tam (200 µM). Data represents mean ± SEM from three replicates. Asterisk denotes significant differences, for all statistical analyses (t-test): *, and ***, indicate P < 0.05,and 0.001, respectively.

**Fig. S6.**
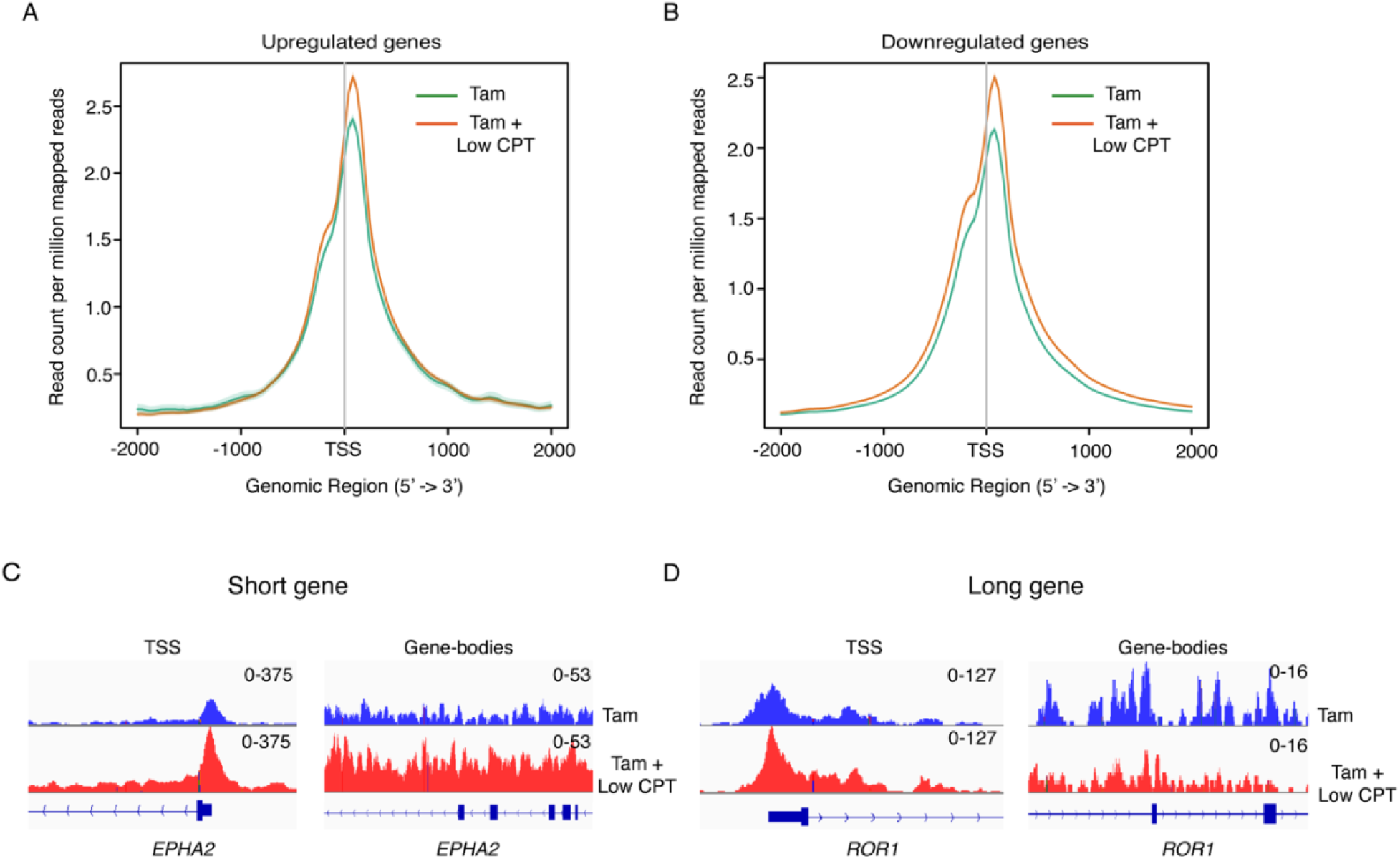
Reduced TOP1 activity enhances RNAPII enrichment and facilitate transcription activation. *(A-B)* Metagene profiles of RNAPII occupancy (read counts per million mapped reads [RPM]) in MCF10A cells expressing lentiviral E-box (-) mCherry constructs after 24 h treatment with indicated combinations of Dox (150 ng/ml), Tam (200 µM) and CPT (100 nM). Metagene analyses of RNAPII occupancy (RPM) are shown at the TSS ± 2000 bp for upregulated genes (A), and downregulated genes (B) under the indicated conditions. (*C-D*) Genome browser tracks showing RNAPII occupancy at the short gene (*EPHA2*) and long gene (ROR1) locus. Signal intensities are displayed for the TSS region (left) and towards the TES (right). Blue traces represent the control [Tam], and red traces represent the Low CPT [Tam + LowCPT] condition.

**Table S1:**
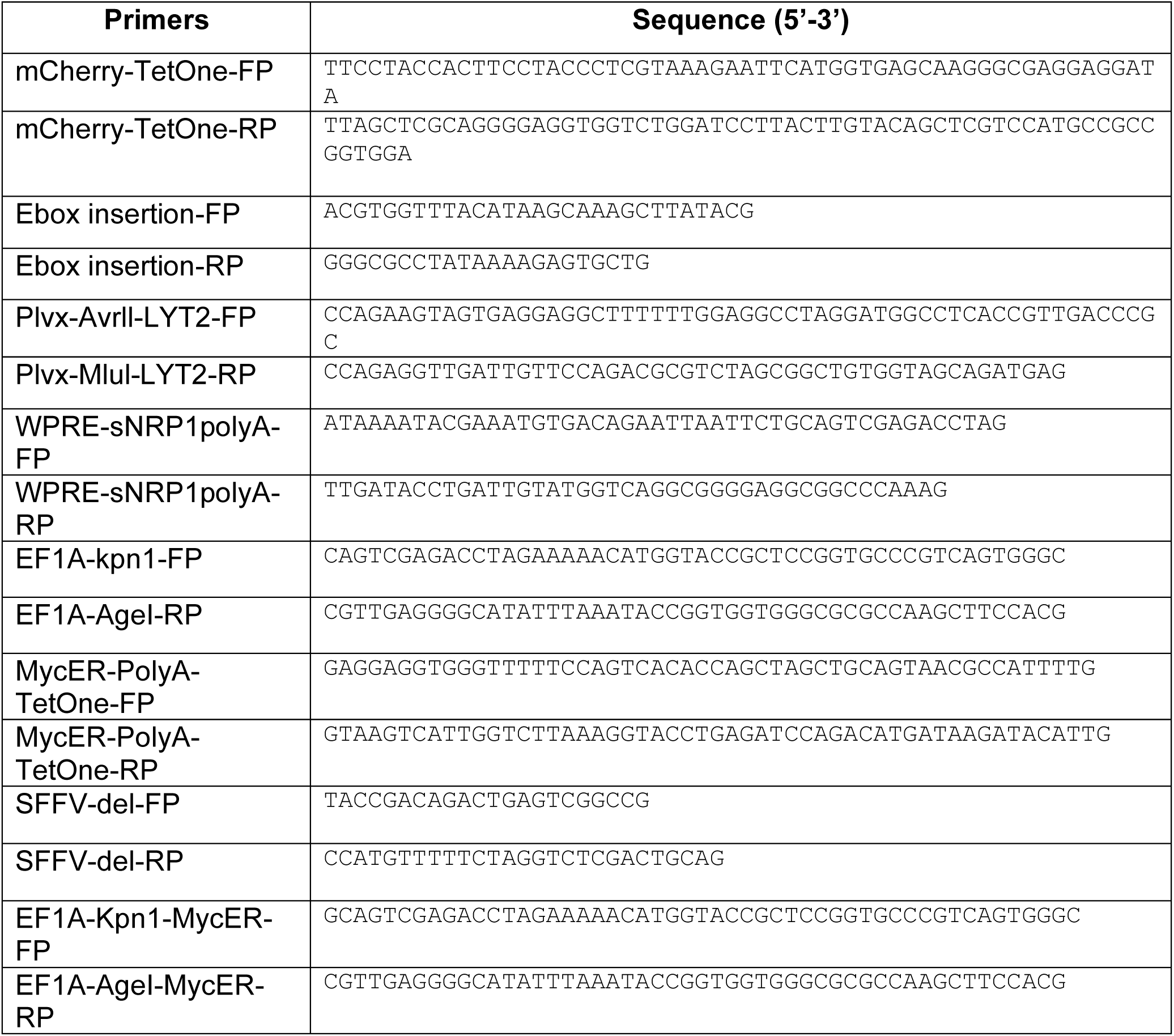
Cloning Primers.

**Table S2:**
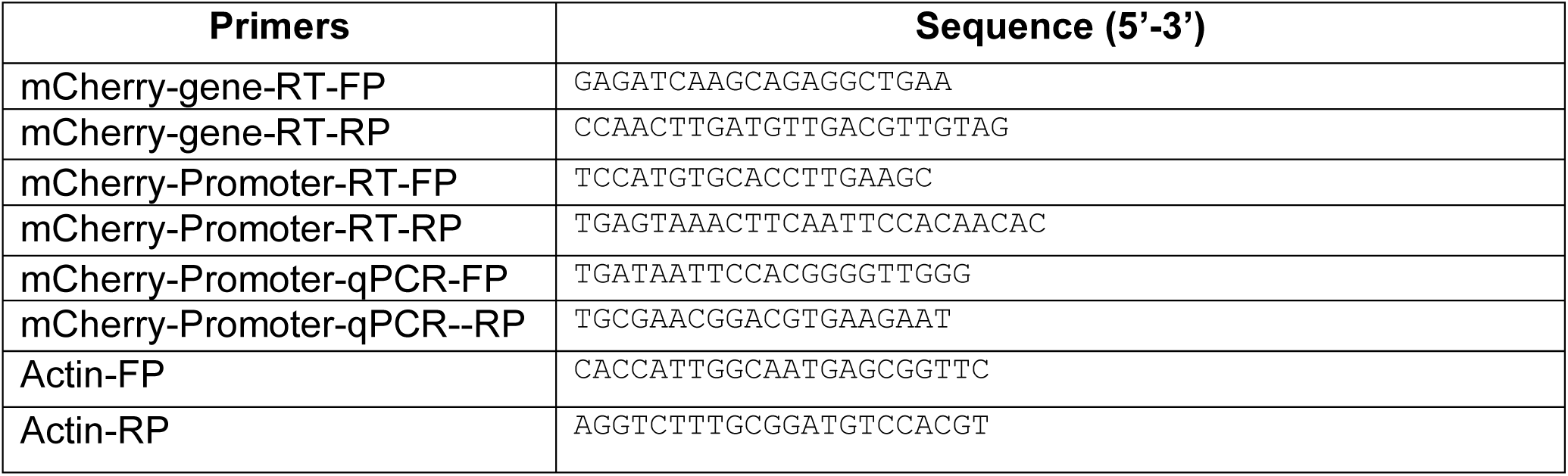
Real-time PCR and qPCR primers.

**Table S3:** Reagents and Source.

| REAGENT or RESOURCE | SOURCE | IDENTIFIER |
| --- | --- | --- |
| **Chemicals** |  |  |
| DMEM, high glucose | Thermo Fisher Scientific | Cat# 11965092 |
| Opti-MEM™ I Reduced Serum Medium | Thermo Fisher Scientific | Cat# 31985062 |
| Corning® 100 mL MEM Nonessential Amino Acids, 100x | Corning | Cat# 25-025-CI |
| Penicillin-Streptomycin (10,000 U/mL) | Gibco | Cat# 15140122 |
| Bovine Serum, heat inactivated, New Zealand origin | Thermo Fisher Scientific | Cat# 26170043 |
| Charcoal/Dextran Stripped Fetal Bovine Serum | GeminiBio | Cat# 100119500 |
| DMEM/F-12 | Invitrogen | Cat# 11330032 |
| Horse Serum, New Zealand origin | gibco | Cat# 16050114 |
| EGF | Peprotech | Cat# AF-100-15 |
| Hydrocortisone | Milipore-Sigma | Cat# H0888 |
| Cholera Toxin | Milipore-Sigma | Cat# C8052 |
| Insulin | Milipore-Sigma | Cat# I1882 |
| DharmaFECT™ 2 Transfection Reagent | Horizon | Cat# T-2002-03 |
| ON-TARGETplus Smartpool Human TOP1 siRNA | Horizon | Cat# L-005278-00-0005 |
| ON-TARGETplus Smartpool Human TOP2A siRNA | Horizon | Cat# L-004239-00-0005 |
| Non-targeting control siRNA | Horizon | Cat# D-001210-02-05 |
| cOmplete™, Mini, EDTA-free Protease Inhibitor Cocktail | Milipore-Sigma | Cat# 11836170001 |
| Tamoxifen | Milipore-Sigma | Cat# T5648-1G |
| ICRF-187 | Selleck Chemicals LLC | Cat# S1222 |
| Etoposide | Milipore-Sigma | Cat# E1383-250MG |
| Bleomycin | Milipore-Sigma | Cat# BP971 |
| Camptothecin | Milipore-Sigma | Cat# C9911-250MG |
| UltraPure™ Agarose | Thermo Fisher Scientific | Cat# 16500500 |
| VECTASHIELD® Antifade Mounting Medium with DAPI | Vector laboratories | Cat# H-1200-10 |
| Pierce™ Protein A/G Magnetic Beads | Thermo Fisher Scientific | Cat# 88803 |
| RNase A, DNase and protease-free (10 mg/mL) | Thermo Fisher Scientific | Cat# EN0531 |
| Sodium deoxycholate monohydrate BioXtra, ≥99.0% (titration) | Milipore-Sigma | Cat# D5670-25G |
| Urea ACS reagent, 99.0-100.5% | Milipore-Sigma | Cat# U5128-5KG |
| DTT 1,4-Dithiothreitol | Milipore-Sigma | Cat# 11583786001 |
| Nonidet™ P 40 Substitute | Milipore-Sigma | Cat# 74385 |
| Ethylenediaminetetraacetic acid disodium salt dihydrate | Milipore-Sigma | Cat# 03685-500G |
| Exonuclease I | New England Biolabs Inc. | Cat# M0293L |
| NuPAGE™ LDS Sample Buffer (4X) | Thermo Fisher Scientific | Cat# NP0007 |
| Pierce™ 16% Formaldehyde (w/v), Methanol-free | Thermo Fisher Scientific | Cat# 28906 |
| 4-12% Bis-Tris NuPAGE protein gel | Thermo Fisher Scientific | Cat# NP0323 |
| SuperSignal™ West Femto Maximum Sensitivity Substrate | Thermo Fisher Scientific | Cat# 34095 |
| Nitrocellulose Membranes, 0.45 µm | Thermo Fisher Scientific | Cat# 77010 |
| Sodium Chloride | Milipore-Sigma | Cat# S9888 |
| Bovine Serum Albumin | Milipore-Sigma | Cat# A7906 |
| Ethyl Alcohol | Milipore-Sigma | Cat# E7023 |
| Magnesium chloride | Milipore-Sigma | Cat# M8266 |
| Direct-zol RNA Miniprep Kits | Zymo-Research | Cat# R2053 |
| ChIP-IT High Sensitivity | Activ Motif | Cat# 53040 |
| SYBR™ Green I Nucleic Acid Gel Stain | Thermo Fisher Scientific | Cat# S7563 |
| Phenol:Chloroform:Isoamyl Alcohol 25:24:1 Saturated with 10 mM Tris, pH 8.0, 1 mM EDTA | Milipore-Sigma | Cat# P2069-400ML |
| Proteinase K, Molecular Biology Grade | New England Biolabs Inc. | Cat# P8107S |
| 4,5',8- trimethylpsoralen | MP Biomedicals | Cat# 154157 |
| 32-CTP 3000 Ci/mmol, 0.25 mCi | Perkin Elmer Lifer Sciences | Cat# BLU008H250UC |
| **Antibodies** |  |  |
| Rabbit monoclonal Anti-Topoisomerase I antibody | Abcam | Cat# ab109374; RRID:AB_10861978 |
| Rabbit monoclonal Recombinant Anti-c-Myc antibody [Y69] | Abcam | Cat# ab32072; RRID:AB_731658 |
| RNA pol II antibody | Active Motif | Cat# 102660 RRID:AB_2687451 |
| normal rabbit IgG antibody | Santa Cruz Biotechnology | Cat# sc-2027; RRID:AB_737197 |
| Goat Anti-Rabbit IgG H&L (HRP) antibody | Abcam | Cat# ab205718;<br>RRID:AB_2819160 |
| mouse monoclonal anti-actin | Cell Signaling Technology | Cat# 3700;<br>RRID:AB_2242334 |
| Alexa Fluor<sup>R</sup> 647 anti-mouse CD8a | BioLegend | Cat# 100724<br>RRID:AB_389326 |
| Rabbit monoclonal Anti-Topoisomerase II alpha antibody | Abcam | Cat# ab52934;<br>RRID:AB_883143 |

